# ReCappable Seq: Comprehensive Determination of Transcription Start Sites derived from all RNA polymerases

**DOI:** 10.1101/696559

**Authors:** Bo Yan, George Tzertzinis, Ira Schildkraut, Laurence Ettwiller

**Affiliations:** New England Biolabs Inc., 240 County Road, Ipswich, MA, 01938, USA

**Author notes:** Both authors contributed equally to this work (Bo Yan, George Tzertzinis). Co-last authors (Ira Schildkraut, Laurence Ettwiller).

## Abstract

Determination of eukaryotic Transcription Start Sites (TSS) has been based on methods that require the cap structure at the 5’ end of transcripts derived from Pol-II RNA polymerase. Consequently, these methods do not reveal TSS derived from the other RNA polymerases which also play critical roles in various cell functions. To address this limitation, we developed ReCappable-seq which comprehensively identifies TSS for both Pol-lI and non-Pol-II transcripts at single-nucleotide resolution. The method relies on specific enzymatic exchange of 5’ m^7^G caps and 5’ triphosphates with a selectable tag. When applied to human transcriptomes, ReCappable-seq identifies Pol-II TSS that are in agreement with orthogonal methods such as CAGE. Additionally, ReCappable-seq reveals a rich landscape of TSS associated with Pol-III transcripts which have not previously been amenable to study at genome-wide scale. Novel TSS from non-Pol-II transcription can be located in the nuclear and mitochondrial genomes. ReCappable-seq interrogates the regulatory landscape of coding and non-coding RNA concurrently and enables the classification of epigenetic profiles associated with Pol-lI and non-Pol-II TSS.

## Introduction

Current widely used methods to characterize transcriptomes such as whole transcriptome shotgun sequencing (RNA-seq) (1) fall short in providing accurate descriptions of transcriptional landmarks such as Transcription Start Sites (TSS), termination sites, and isoform composition. The identification of TSS is essential to study gene regulation because it permits the association between RNA transcription and the underlying genomic landmarks such as promoters and histone marks. Alternative TSS have been detected in more than 50% of human genes (2) driving most of the transcript isoforms differences across tissues (3). Genomic positions of alternative TSS are often found within other isoforms of the same gene, confounding their detection using conventional RNA-seq.

Currently methods for TSS determination such as Cap Analysis of Gene Expression (CAGE)(4,5), NanoCAGE (6), and Oligo-capping (7) are limited to identifying 7mG capped Pol-II derived TSS. These methods entirely exclude the large number of TSS derived from eukaryotic RNA polymerase I (Pol-I), RNA polymerase III (Pol-III), and mitochondrial RNA polymerase (POLRMT) which produce uncapped non-coding RNA. These uncapped primary transcripts display a 5’ triphosphate identical to the 5’ end of prokaryotic primary transcripts. With the growing body of literature highlighting the key role of both Pol III and Pol II transcribed non-coding RNA in regulating biological processes and diseases (8, 9, 10) a method that comprehensively identifies TSS for all eukaryotic RNA polymerases would be consequential.

We have previously developed Cappable-seq to identify TSS in prokaryotic species (11). Cappable-seq is based on the ability of the Vaccinia Capping Enzyme (VCE) to add a biotinylated guanosine to 5’ di- or triphosphorylated RNA ends and streptavidin enrichment of those fragments. Building on Cappable-seq, we have developed ReCappable-seq to also capture 7-methyl G-capped transcripts derived from Pol-II RNA polymerase. To achieve this, we took advantage of the property of the yeast scavenger decapping enzyme (yDcpS) to convert capped RNA (12) into di-phosphorylated RNA that can be “re-capped” by the vaccinia capping enzyme (13). Thus, ReCappable-seq enables the identification of TSS for RNA transcripts derived from all RNA polymerases.

The comparison of two datasets, one derived from RNA treated with calf intestinal alkaline phosphatase (CIP) and the other not, permits the discrimination between capped 5’ ends and triphosphorylated 5’ ends. Each TSS can be inferred as derived either from Pol-II or non-Pol-II polymerases, because Pol-II transcripts are capped and not depleted with CIP while non-Pol-II transcripts are triphosphorylated and depleted with CIP.

We applied ReCappable-seq to the transcriptome of the A549 human cancer cell line and identified 33,468 and 5,269 Pol-II and non-Pol-II TSS respectively. Among the non-Pol-II TSS, we detected the known Pol-I TSS located upstream of the annotated 45S ribosomal RNA locus and 2 out of the 3 known mitochondrial TSS. We also identified TSS at over 80% of the Pol-III occupied sites detected by ChIP-seq in HeLa cells (14). Importantly, we identified over one thousand of novel non-Pol-II TSS revealing a rich landscape of the non-coding primary transcriptome. Pol-II TSS identified by ReCappable-seq are in substantial agreement with CAGE with 97% of ReCappable-seq TSS located within 1bp of a CAGE signal. Furthermore we demonstrate that ReCappable-seq can identify TSS from highly degraded RNA samples. ReCappable-seq provides an accessible, affordable method to comprehensively follow the primary transcriptome not only of Pol-II-derived m^7^G capped RNA but also of the primary transcripts derived from the other RNA polymerases.

## Results

### Preparation of ReCappable-seq libraries

The principle of ReCappable-seq (shown schematically in Figure 1a) relies on the tagging of all primary transcripts with biotin. RNA is subjected to decapping with yDcpS which acts on capped transcripts originating from Pol-II transcription. yDcpS hydrolyzes the phosphodiester bond between the gamma and beta phosphates of the 7mG-cap (7mGppp-RNA) leaving a diphosphate end (13). Importantly, cap0 and cap1 as well as m7Gpppm6A and m7Gpppm6Am are all substrates for yDcpS (13). Subsequently, the RNA is capped with a biotin-modified GTP analog (3’-desthiobiotin-GTP) using VCE. At this step, the decapped diphosphate RNA (pp-RNA) product of yDcpS and the 5’ triphosphorylated RNA (ppp-RNA) originating from Pol-I, Pol-III, and POLRMT (non-Pol-II) transcription are capped with the biotin-modified analog. This biotinylation step allows enrichment of all primary transcripts on a streptavidin matrix (Figure 1a). Differentiation of Pol-II from the non-Pol-II transcripts is accomplished by sequencing a second library constructed with RNA treated with CIP prior to the yDcpS treatment, in order to remove the 5’ triphosphate from non-Pol-II transcripts (Figure 1a). This treatment prevents the non-Pol-II transcripts from being capped with biotin and enriched. Comparison of these two libraries allows the distinction of capped transcripts from 5’ triphosphate transcripts. The enriched RNA from both libraries is subsequently decapped with RppH to generate 5’ monophosphate ends which are captured in a ligation-based library.

**Figure 1:**
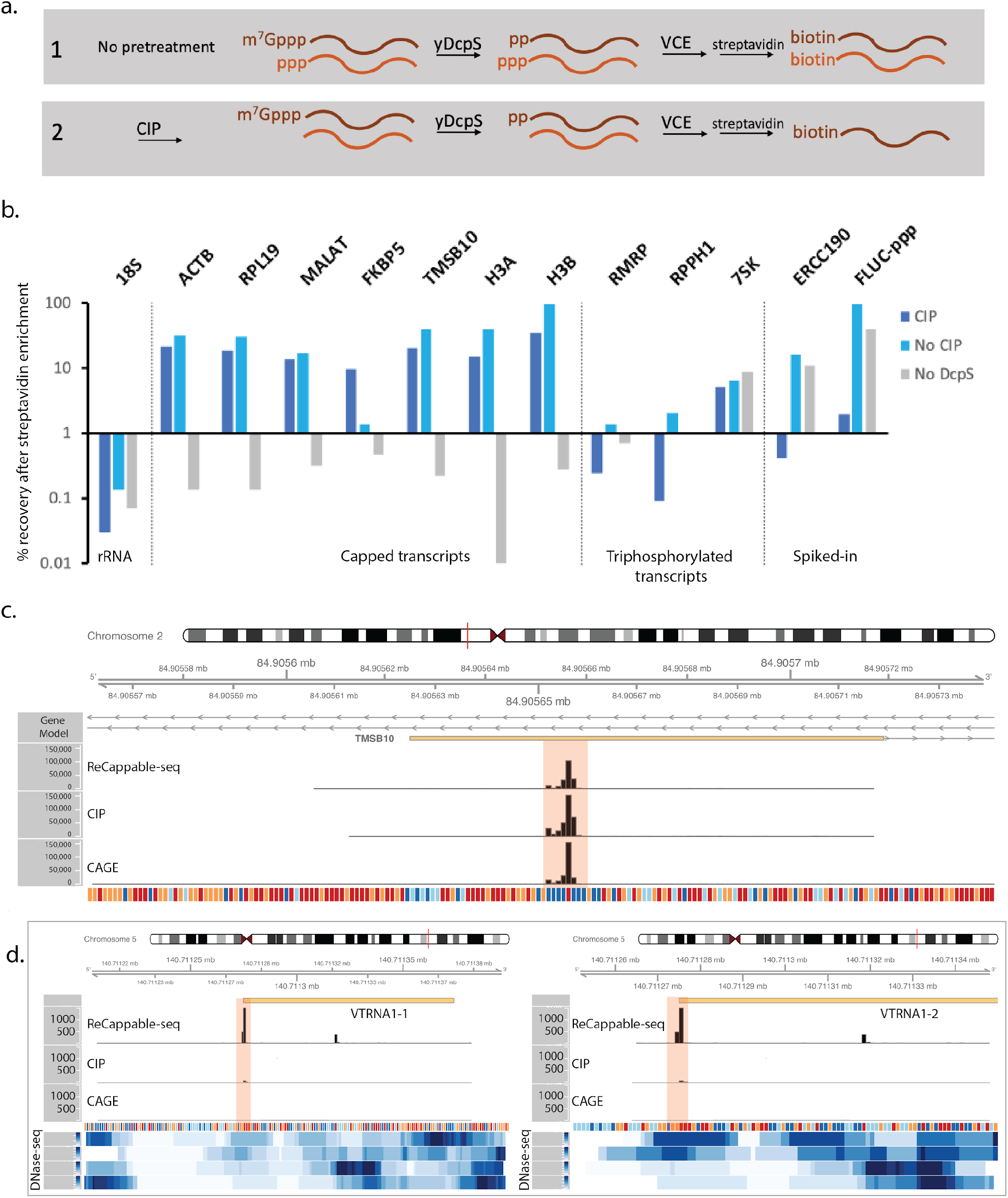
ReCappable-seq. a. Principle of ReCappable-seq. 1. RNA is subjected to decapping with yDcpS which acts on capped transcripts originating from Pol-II transcription. Subsequently, the RNA is capped with a biotin-modified GTP analog (3’-desthiobiotin-GTP) using VCE. This biotinylation step allows enrichment of all primary transcripts on a streptavidin matrix. 2. Differentiation of Pol-II from the non-Pol-II transcripts is accomplished by sequencing a second library constructed with RNA treated with CIP prior to the yDcpS treatment, in order to remove the 5’ triphosphate from non-Pol-II transcripts.b. RT-qPCR assay measuring the recovery after streptavidin enrichment of various classes of transcripts such as 18S rRNA as an example of a processed transcript (rRNA), ACTB, RPL19, MALAT (MALAT1), FKBP5, TMSB10, H3H (HIST1H3H) and H3B (HIST1H3B) as examples of capped transcripts, RMRP, RPPH1 and 7SK (RN7SK) as examples of Pol III transcripts (with 7SK having a 5’ methylated triphosphate and therefore resistant to CIP treatment, see main text) and ERCC190 and FLUC-ppp as examples of spiked-in *in vitro* transcripts with a defined triphosphorylated 5’ end. c. Example of a Pol-II TSS in the TMSB10 locus: the same positions (shaded in pink) are found in the CAGE dataset. CIP treatment intensifies the signal, consistent with a Pol-II TSS. d. Example of Pol-III TSS corresponding to the start of two vault RNAs (vault RNA 1-1 and vault RNA 1-2) located on Chr.5. The positions (shaded in pink) are missing in the CAGE dataset. CIP treatment reduces the signal, consistent with non-Pol-II TSS. In c. and d. the tracks correspond to ReCappable-seq, CIP-Recappable-seq, CAGE, and RNA-seq (A549 rRNA-depleted RNA-seq) read coverage. The four bottom tracks correspond to read density from public ENCODE DNase-seq from A549 cells (ENCFF473YHH, ENCFF809KIH, ENCFF821UUL, ENCFF961WXW).

### Validation of the ReCappable-seq principle by RT-qPCR

In order to demonstrate that primary transcripts bearing either a 5’ cap or a 5’ triphosphate can be specifically captured from total RNA, we subjected total RNA from A549 cells to the enzymatic and streptavidin steps described in Figure 1a and quantified specific transcripts using RT-qPCR. In order to demonstrate the requirement of the decapping step we also processed samples for which the yDcpS step was omitted.

The results showed substantial recovery of both Pol-II and Pol-III primary transcripts after streptavidin enrichment, while 18S ribosomal RNA, a non-capped, non-triphosphorylated transcript, was significantly depleted (Figure 1b). As expected, when the decapping step was omitted, capped transcripts were depleted whereas triphosphorylated Pol-III transcripts were recovered (Figure 1b). Conversely, when RNA was CIP treated, capped transcripts were recovered whereas triphosphorylated Pol-III transcripts were depleted (Figure 1b). An exception is 7SK, known to possess a monomethyl gamma-phosphate at its 5’ end, which is resistant to CIP (24).

These results demonstrate that the yDcpS decapping and VCE recapping steps specifically allow the recovery of capped transcripts and that CIP treatment enables the distinction of m^7^G capped RNA from triphosphorylated RNAs (Figure 1b).

### Genome-wide identification of TSS using ReCappable-seq

We applied ReCappable-seq to total RNA isolated from the human cell line A549. Two samples (untreated and pretreated with CIP) were processed using the steps outlined in Figure 1a. The RNA was then ligated with adaptors to prepare libraries suitable for short read high throughput sequencing as previously described in (11) (See Materials and Methods for protocol details). This strategy results in the genome-wide identification of TSS derived from all RNA polymerases at single-nucleotide resolution.

In order to evaluate the reproducibility of ReCappable-seq, we utilized technical replicates yielding approximately 32 million single-end Illumina reads per library (Supplementary Table). In addition, for both the CIP-treated and untreated samples, unenriched libraries were constructed for each from an aliquot taken prior to the streptavidin enrichment step. Reads were mapped to the human genome using STAR with the ENCODE default parameters (Materials and Methods)(15).

In parallel, we used RNA from the same A549 RNA preparation and performed CAGE for comparison. Analysis of the ReCappable-seq technical replicates at single-nucleotide resolution reveals a high correlation (Pearson corr.=0.96, P-value < 2.2e-16) between replicates (Supplementary Figure 1a), demonstrating high reproducibility of the technique. ReCappable-seq replicates were combined and downsampled to 63 million mappable reads for subsequent analyses.

We evaluated the specificity of ReCappable-seq for primary transcripts (5’ m^7^G capped or 5’ triphosphorylated ends) versus non-primary transcripts, using the fraction of reads mapping to rRNA as a surrogate for non-primary transcripts. rRNAs are formed by the processing of a single pre-rRNA 45S transcript to form the mature 18S, 5.8S and 28S rRNAs (16). rRNAs account for the vast majority of the RNA mass in the cell and because they are processed, they are expected to be depleted in ReCappable-seq libraries. Accordingly, the percentage of rRNA mapped reads drops from about 70% in the unenriched control libraries to 3-4% in the ReCappable-seq libraries (Supplementary Figure 2a, Supplementary Table), highlighting the specificity of ReCappable-seq and its efficiency in removing transcripts with processed or degraded 5’ ends.

Mapped reads are found near the 5’ end of both protein-coding transcripts (Supplementary Figure 2b) as well as non-coding transcripts known to be transcribed by Pol-III (see Figure 1c and 1d for examples) suggesting high specificity for all types of TSS.To further investigate the specificity of ReCappable-seq for primary 5’ ends, we tested ReCappable-seq on pre-fragmented RNA samples (RIN number <3) prepared by magnesium ion-mediated fragmentation to simulate naturally occurring RNA degradation in biological samples. RNA degradation has been proven to be challenging for the determination of TSS because the majority of the 5’ ends are generated from fragments and do not correspond to TSS.

The profile of mapped reads demonstrates a high correlation between both pre-fragmented replicates (Pearson corr=0.97, Supplementary Figure 1c) and reasonable correlation between pre-fragmented and intact starting material (Pearson corr=0.76, Supplementary Figure 2c). Furthermore reads from pre-fragmented material predominantly map to the start of annotated genes consistent with the positioning of TSS (Supplementary Figure 2b and d). Importantly, reads mapping to processed rRNA drop from 65% in the unenriched control libraries to ∼1.3% in the ReCappable-seq libraries derived from pre-fragmented RNA (Supplementary Figure 2a). Thus, ReCappable-seq is not affected by the large excess of uncapped 5’ ends resulting from fragmentation, and genuine primary 5’ ends were predominant in the ReCappable-seq libraries. Together, these results show that ReCappable-seq performs well on intact and fragmented RNA, adding further support to its high specificity for identifying 5’ ends of primary transcripts.

### Classification of TSS into Pol-II and non-Pol-II TSS

Candidate TSS were identified and quantified at single-nucleotide resolution using a TSS analysis pipeline (Materials and Methods). In short, candidate TSS were defined as single nucleotide positions in the genome where the number of 5’ ends of the reads mapping to those is above 1 TPM (TSS -tags per million mapped reads). From 63 million primary mappable reads, a total of 42,988 candidate TSS were identified genome-wide.

To evaluate the specificity of ReCappable-seq in capturing capped and triphosphate RNAs, an enrichment score was calculated for each candidate TSS by dividing the TPM in the ReCappable-seq libraries with the TPM in the unenriched control library (Ct)(Figure 2a). Candidate TSS depleted in the ReCappable-seq library (ratio less than 1) are considered false positives (Figure 2b quadrants I and III). The enriched candidate TSS (ratio equal to or greater than 1) are considered high confidence TSS (Figure 2b quadrants II and IV). 38,737 of the 42,988 candidate TSS were high confidence indicating that the majority (90.1%) of the TSS identified by ReCappable-seq are true positives. Unless otherwise stated below, the term TSS refers to the high confidence TSS.

**Figure 2:**
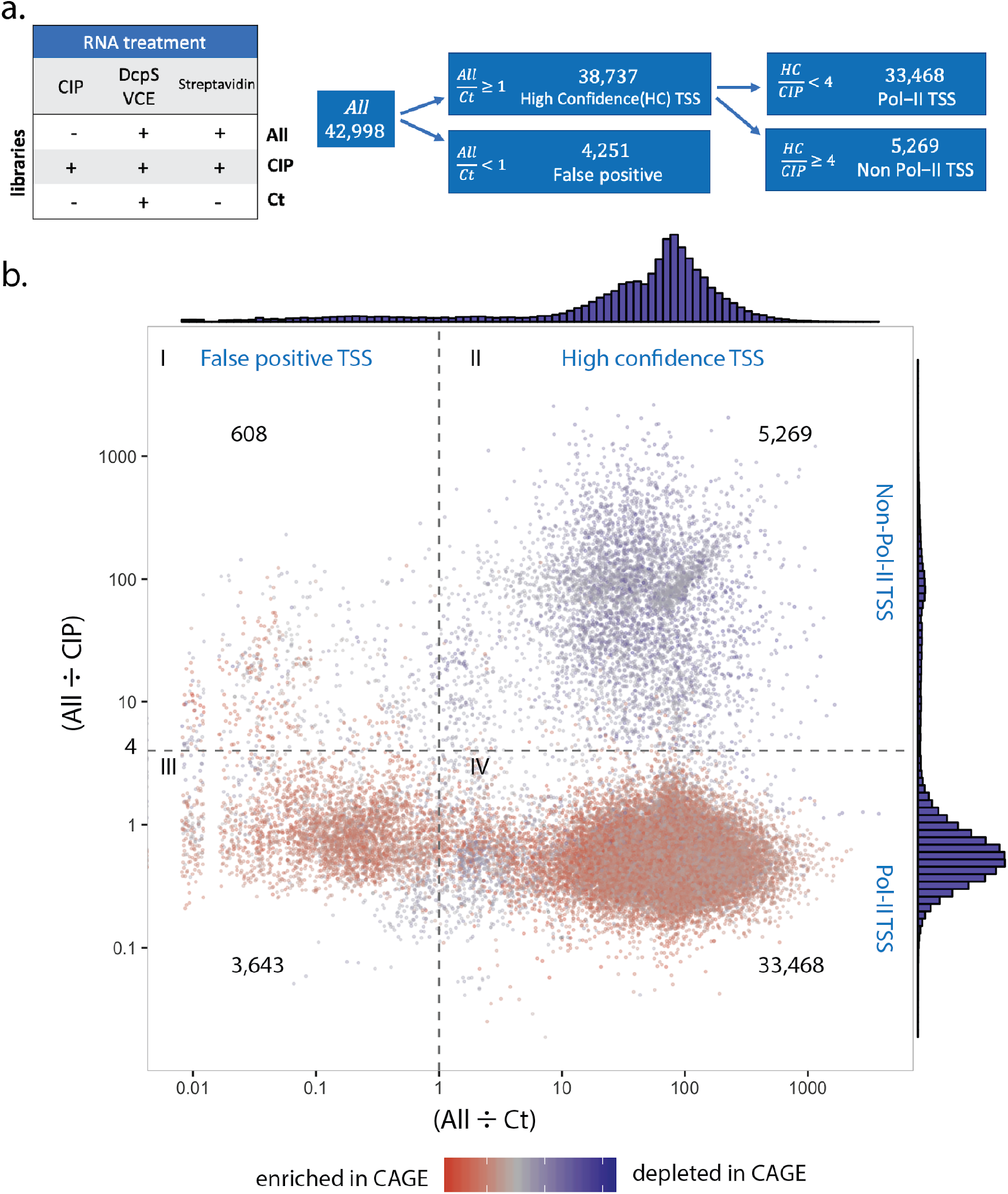
a. Summary of sequencing experiments: Experiments were performed resulting in 3 dataset types: ReCappable-seq (ALL), CIP treated ReCappable-seq (CIP), and unenriched control for which the streptavidin step has been omitted (Ct). ALL corresponds to the ReCappable-seq dataset without CIP treatment and defines 42,988 candidate TSS. Comparison of ALL with Ct enables the definition of 38,737 high confidence (HC) TSS. Comparison of HC with CIP enables the definition of high confidence Pol-II TSS (33,468) and the high confidence non-Pol-II TSS (5,269). b. 42,988 candidate TSS positions distributed according to the TPM ratio between ALL and Ct (x-axis) and the TPM ratio between ALL and CIP (y-axis). Dotted lines define 4 quadrants as follows: ALL÷CIP less than 4 (Pol-II TSS), ALL÷CIP above or equal to 4 (non-Pol-II TSS), ALL÷Ct above or equal to 1 (high confidence TSS), and ALL÷Ct less than 1 (false positive TSS). Colors denote the ratio between ALL and CAGE with red (enriched in CAGE relative to ReCappable-seq), blue (depleted in CAGE relative to ReCappable-seq).

To enable the discrimination of TSS derived from capped transcripts from those derived from triphosphate transcripts, we used the data from RNA pretreated with CIP prior to performing ReCappable-seq (Figure 1a, Supplementary Figure 1b). Comparison of the non-CIP ReCappable-seq dataset with the CIP treated dataset reveals two populations composed of 33,468 CIP resistant TSS consistent with Pol-II transcripts and 5,269 CIP sensitive TSS consistent with non-Pol-II transcripts (Figure 2b quadrants II and IV respectively).

### TSS consistent with Pol-II transcripts

We validated Pol-II consistent TSS from ReCappable-seq using data from orthogonal methods known to mark TSS, such as CAGE and publically available data for DNase-seq, chromatin immunoprecipitation (ChIP-seq) for Pol-II and manually curated human gene annotation. For the CAGE datasets, we used datasets derived from the same A549 RNA preparation used for ReCappable-seq.

We assessed precision and sensitivity of ReCappable-seq following the published protocol as described in (17). We compared ReCappable-seq with CAGE and found that 97% of the ReCappable-seq TSS have a CAGE signal within a window of +/-1 nucleotide (Figure 3a). When compared against gene annotation and DNase-seq we found precision of 81% and 90% respectively (Figure 3a). When CAGE is compared to DNase-seq sites and gene annotation, the accuracy is similar to ReCappable-seq. CAGE shows higher sensitivity presumably due to the fact that ReCappable-seq data have less sequence depth for Pol-II TSS because they include non-Pol-II TSS accounting for ∼50% of the sequencing reads (Figure 3b).

**Figure 3:**
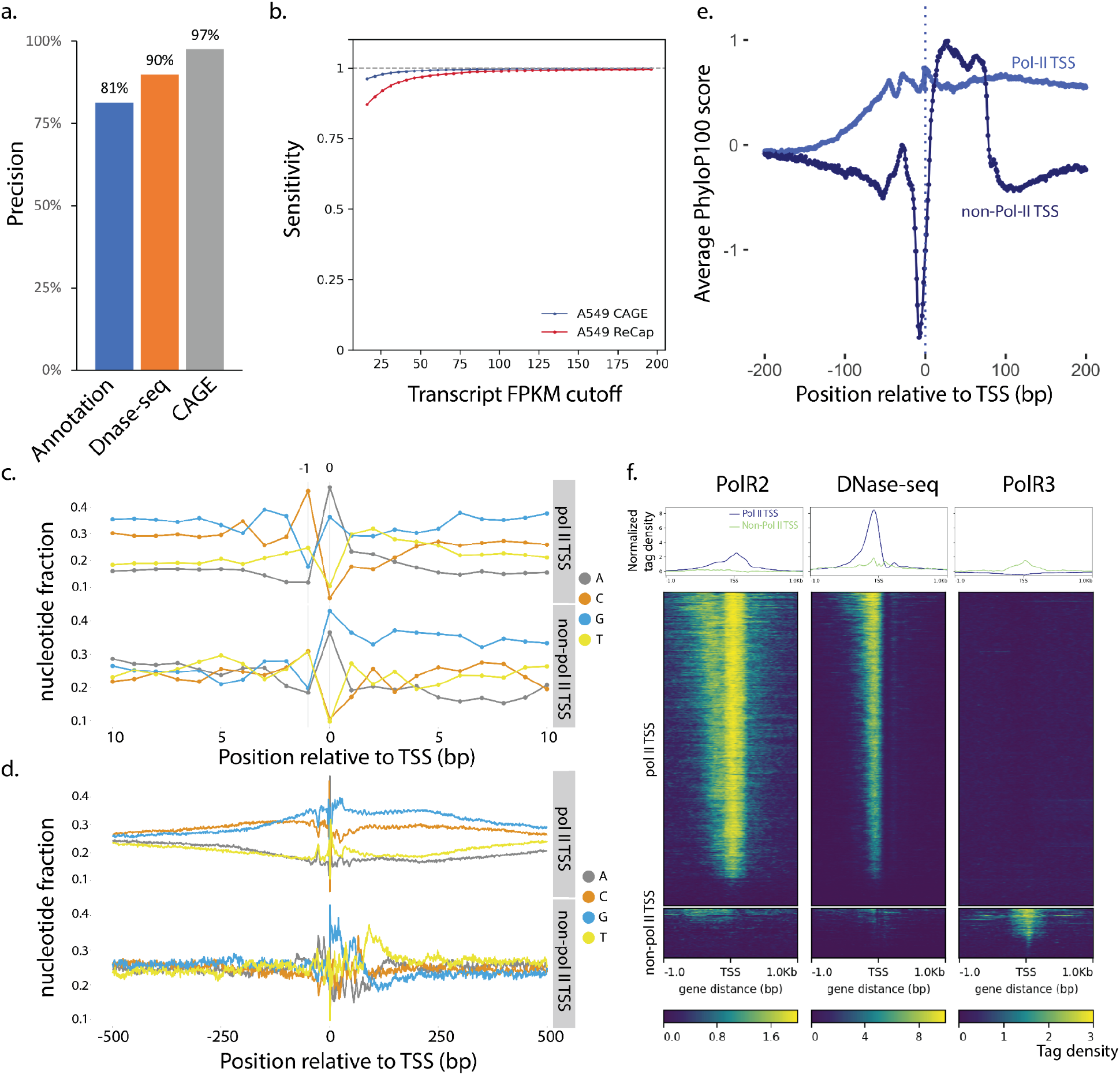
TSS Characterization: a. Precision (TP/(TP+FP))*100 of ReCappable-seq with TP = true positive, FP = False positive (see Material and Methods). b. Sensitivity (TP/(TP+FN)) of ReCappable-seq with TP = true positive (ReCappable-seq TSS with UCSC annotation) and FN = False negative (TSS in UCSC annotation but not detected by ReCappable-seq) (see Material and Methods) c. Nucleotide composition in the 20 bp flanking region for Pol-II TSS (upper) and non-Pol-II TSS (lower). d. Same as c. for 1kb flanking region. 83.7% and 79.5% of Pol-II and non-Pol-II TSS respectively start with A or G. e. Conservation profiles using PhyloP basewise conservation score (32) derived from Multiz alignment (33) of 100 vertebrate species around Pol-II TSS (black) and non-Pol-II TSS (orange). f. Profiles and heatmaps of Pol-II (left panel) and Pol-III (right panel) ChIP-seq and DNase-seq (middle panel) at Pol-II TSS and non-Pol-II TSS. The Pol-II ChIP-seq and DNAse-seq data have been downloaded from the ENCODE website (34), the Pol-III ChIP-seq data have been obtained from (14).

### Comparison between Pol-II and non-Pol-II TSS

We analyzed the nucleotide composition for Pol-II and non-Pol-II TSS and found that 83.7% and 79.5% respectively of the +1 nucleotides are A or G and 70.6% and 61.7% respectively of the -1 position are C or T (Figure 3c, d) revealing the canonical -1[C or T] +1[G or A] motif typical for TSS for both Pol-II and non-Pol-II TSS.

Limiting TSS to uniquely mapped reads, we interrogated the conservation profiles around Pol-II and non-Pol-II TSS. We found a peak of conservation for Pol-II TSS consistent with the fact that these sites are functional and therefore under selection (Figure 3e). Interestingly for non-Pol-II TSS, the conservation profile is very different from the Pol-II TSS indicating a very different evolutionary conservation for the two classes of transcripts (Figure 3e).

Next, using data from ENCODE Pol-II ChIP-seq from A549 cells and Pol-III ChIP-seq from HeLa cells (14) (the only Pol-III ChIP-seq data publicly available), we interrogated the binding profiles of both RNA polymerases relative to ReCappable-seq derived Pol-II and non-Pol-II TSS. Because the Pol-III ChIP-seq data are from a different cell line, concurrence of Pol-III polymerase binding sites with TSS may only be found for commonly expressed genes between HeLa and A549. Consistent with the origin of Pol-II and non-Pol-II TSS, we found a higher density of Pol-II ChIP tags (14) around Pol-II TSS (Figure 3f) and a higher density of Pol-III ChIP tags around non-Pol-II TSS (Figure 3f). Interestingly, the Pol-II TSS also show a depletion of Pol-III ChIP-seq tags relative to the flanking regions suggesting that genomic regions at active Pol-II TSS are devoid of bound Pol-III.

It has been shown that the chromatin landscape of Pol-III-transcribed genes in many ways resembles that of Pol-II-transcribed genes, although there are clear differences (18, 19). Substantial differences in chromatin structure of Pol-II and Pol-III promoters have been reported (20). With the ability to identify and classify Pol-II and non-Pol-II consistent TSS, we studied the differential positioning of chromatin marks and DNA-interacting proteins relative to both TSS classes. For this, we compared ReCappable-seq TSS with published chromatin immunoprecipitation (ChIP-seq) from ENCODE data performed on A549 cells to identify interesting differential chromatin marks associated with Pol-II and non-Pol-II consistent TSS.

Most of the chromatin landscape around TSS of non-Pol-II transcribed genes resembles that of Pol-II transcribed genes in agreement with the literature (18, 19) (Supplementary Figure 5). Nonetheless, we find examples of specific transcription factor binding profiles at non-Pol-II transcribed genes that have not been previously reported to be associated with the regulation of non-Pol-II genes (Supplementary Text 1, Supplementary Figure 5). For example, SREBP2 (SREBF2) a ubiquitously expressed transcription factor known to control cholesterol homeostasis (21) is binding almost exclusively at non-Pol-II TSS. This result strongly suggests a role of SREBP2 in regulating non-Pol-II genes.

Distinct DNase I accessibility profiles can be observed at Pol-II versus non-Pol-II TSS. Pol-II transcripts show maximum DNase I accessibility a few nucleotides upstream of their TSS (Figure 3f) consistent with nucleosome depletion at Pol-II TSS (22). In contrast, non-Pol-II transcripts show minimal DNase I accessibility at TSS (Figure 3f). Distinctive nucleosome positioning has been previously observed for promoters of Pol-II and Pol-III RNA polymerases (20). Our results further extend the distinctive accessibility of DNA at TSS of Pol-II and non-Pol-II transcripts and highlight a distinct chromatin landscape for Pol-II and non-Pol-II TSS.

Pol-II TSS are found for 7,186 annotated genes. Consistent with the function of Pol-II polymerase, the majority (89%) of ReCappable-seq Pol-II TSS are found either upstream or within protein-coding genes (Figure 4a).

**Figure 4:**
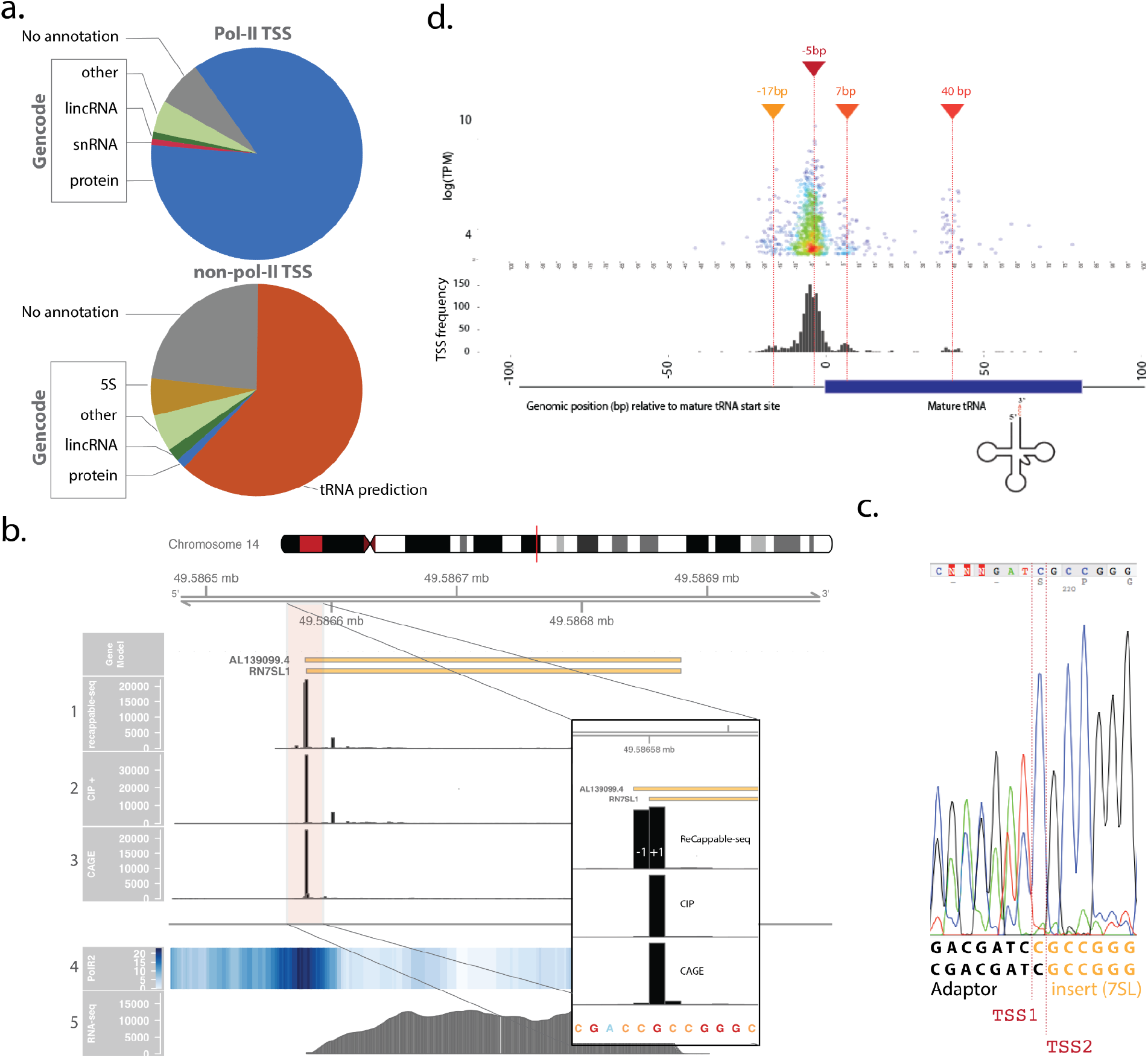
a. Pie charts representing the proportion of TSS associated with GENCODE genes, predicted tRNAs (orange) or not associated with any annotation (grey) for Pol-II (upper chart) and non-Pol-II (lower chart). b. 7SL (RN7SL1) locus showing the distribution of the 5’ end of mapped reads for Recappable-seq (track 1), CIP treated Recappable-seq (track 2) CAGE (track 3), Pol-II ChIP-seq (track 4, read density) and RNA-seq reads (track 5). The floating panel represents a close up of the 5’ end of the RN7SL1 with two TSS at -1 and +1 of the annotated RN7SL1. The 5’ mapped end of the reads is shown for Recappable-seq (track 1), CIP treated Recappable-seq (track 2), and CAGE (track 3) to mark the TSS positions. c. Validation of the two TSS identified for 7SL using RACE. The amplified fragments were directly sequenced using sanger sequencing with a primer located in the 7SL gene (Material and Methods). The sequencing trace reveals two products ligated to the RACE adaptor resulting from two alternative transcript starts corresponding to TSS1 and TSS2. d. Non-Pol-II TSS flanking tRNA annotations. The top panel visualizes individual non-Pol-II TSS relative to the 5’ end of the annotated mature tRNA (in bp) as a function of the TPM. The bottom panel represents the distribution of the non-Pol-II TSS relative to the start of the annotated tRNA starts (in bp).

### Non-Pol-II TSS

Non-Pol-II TSS account for 5,269 positions. In addition to the correlation of the TSS to Pol-III ChIP-seq (Figure 3f), we validated non-Pol-II consistent TSS using 5’ RACE (Supplementary Figure 6). The 5’ RACE results for two uniquely mapped annotated Pol III transcribed genes (RMRP and RPPH1) confirmed that the start positions were the same as the major TSS detected from the ReCappable-seq data.

The 5,269 non-Pol-II TSS (TPM>=1) represent about 50% of the total reads despite representing only 14% of all TSS positions, consistent with the fact that Pol-I and Pol-III transcripts are abundant. Of these non-Pol-II TSS, 4,032 are detected within or upstream of 758 genes of which 314 are annotated as Pol-III transcribed genes such as 5SRNA, Vault RNAs, RMRP, RPPH1, 7SL, YRNA. The other 444 are located at predicted tRNA genes (30) (Figure 4a, Supplementary Figure 3). The remaining 1237 detected TSS correspond to neither GENCODE annotation nor tRNA predictions by tRNAscan, and are distributed in ∼700 loci (Material and methods).

Around one-third of the non-Pol-II TSS (34%) are derived from reads mapping to multiple positions in the genome, consistent with the role of Pol-III polymerase in transcribing repeat elements and multiple-copy genes (18) such as 5S, 7SK and 7SL (23). We, therefore, proceeded with a genome-wide investigation of non-Pol-II TSS with respect to such genes and located them upstream or at the starts of 5S, 7SL, tRNA and U6 genes, and HY and MIR repeats (Supplementary Figure 4a). This result is in agreement with literature that has assigned transcription of these genes to Pol-III (18). Interestingly, a closer look at the 7SL gene identifies 2 strong TSS at +1 and -1 bp relative to the annotated gene start (Figure 4b). The -1 TSS is mostly eliminated by CIP treatment consistent with the triphosphorylated nature of the 5’ end. Conversely, the +1 TSS is not affected by CIP treatment, consistent with a possible canonical cap structure at the 5’ end of 7SL. We experimentally confirmed the presence of both TSS using 5’ RACE followed by Sanger sequencing (Figure 4c, Materials and Methods). The occurrence of a canonical cap structure at +1 of 7SL would sharply contrast with the body of literature that describes 7SL genes as being transcribed by Pol-III (23). Alternatively, the 5’ end of the +1 TSS form may not be accessible to CIP treatment leading to its incorrect assignment to Pol-II. However, in further support of a Pol-II TSS, we also observe a CAGE signal.

#### 7SK

The TSS identified at the start of 7SK genes appear resistant to CIP treatment and as such were classified as Pol-II TSS (Supplementary Figure 4a). Consistently, RT-qPCR results showed that neither CIP nor yDcpS treatment affect the recovery of 7SK upon streptavidin enrichment (Figure 1b). It has previously been shown that the 5’ end of the 7SK transcript contains a monomethyl gamma-phosphate on the 5’ triphosphate (24) conferring resistance to alkaline phosphatase (25). We have confirmed this experimentally (Supplementary Figure 4b, Materials and Methods). Furthermore, we show that VCE can remove the gamma-methylated phosphate indicating that the RNA triphosphatase activity of VCE is not blocked. This suggests that such 5’ ends are subject to *in vitro* capping (Supplementary Methods, Supplementary Figure 4b). Together, these results explain why 7SK TSS are classified as non-Pol-II (CIP resistant and not dependent on yDcpS decapping.)

#### U6

The same result should have been obtained for the U6 small nuclear RNA which is also known to possess a gamma-methyl triphosphorylated 5’ end (26, 27). However, we found no TSS signal from the U6 genes at the annotated start, instead, we found CIP sensitive TSS around 20 bp upstream (Supplementary Figure 4a), consistent with a non-methylated canonical triphosphate RNA 5’ end originating at these positions. To rule out an uncharacteristic cell-line specific expression of U6 in A549, we also performed ReCappable-seq on human brain RNA.

Because U6 genes are highly repeated, we modified the parameters for multiple mapping allowing reads to map to more loci than the default ENCODE mapping parameters. TSS signals at the 5’ end of U6 annotated genes can be found for both the A549 and human brain (see Supplementary text 1 and Supplementary Figure 4c). Nonetheless, TSS at annotated U6 starts are derived from very few reads in both samples, inconsistent with the abundance of U6 (28).

To further explore the U6 TSS findings, we performed 5’ RACE on total A549 RNA using a complementary internal U6 primer (Materials and Methods). After amplification and sequencing of the resulting cDNA, we found evidence for both the annotated and the upstream TSS (Supplementary Figure 4c). We did not recover a substantial amount of the expected U6 transcripts, either because of secondary structure or some other unknown feature of the RNA that prevents capture by our enrichment and cloning protocol.

#### tRNAs

Pre-tRNAs have been notoriously difficult to study due to the rapid processing of primary transcripts relative to the exceptional stability of the mature tRNA which consequently accumulates in the cell (29). With the ability of ReCappable-seq to capture primary transcripts, we are now in a unique position to interrogate the TSS landscape of tRNAs. Using annotated tRNAs from GtRNAdb (30), we found 3,245 of the non-Pol-II TSS are located upstream or within 444 tRNA genes representing the largest class of non-Pol-II TSS found by ReCappable-seq (Figure 4d and Supplementary Figure 3). Notably tRNA-Leu-TAG-1-1 is the highest expressed tRNA (TPM 53181) accounting for approximately 10% of the total non-Pol-II reads in A549 cells.

TSS positions relative to the tRNA annotation highlight a large number of TSS approximately 5 bp upstream of mature tRNA 5’ ends (Figure 4d) consistent with previous work using *in vitro* transcription which identified TSS located mostly 10 to 2 bp upstream of mature plant tRNAs (31). Additionally, we found 3 novel TSS clusters at around -17, +7, and +40 bp from the mature tRNA 5’ end (Figure 4d). Further refinement relative to the type of tRNA reveals that downstream TSS are mostly found in leu tRNA and lys tRNA (Supplementary Figure 7).

#### Identification of the human mitochondrial TSS

The property of ReCappable-seq to capture 5’ di- and triphosphorylated transcripts offers the unique ability to interrogate the TSS landscape of the mitochondrial genome (Supplementary Figure 8a). The entire mitochondrial genome is known to be transcribed from both strands as long polycistronic transcripts (35). The light strand promoter (LSP) controls the transcription of eight of the tRNAs and the MT-ND6 gene. On the heavy strand, a two-promoter system (HSP1 and HSP2) has historically been proposed to explain the higher abundance of the rRNAs. However, the two promoter model remains controversial as more recent experiments (36, 37) suggest that heavy strand transcription is under the control of a single promoter (HSP1) and that the difference in abundance may be a consequence of differential turnover (35).

In agreement with the literature, ReCappable-seq identifies LSP and HSP1 TSS at nucleotide resolution: Both TSS have a strong major peak and a few minor peaks indicative of an imprecise start of transcription (Supplementary Figure 8b and c). Interestingly, most of the transcripts starting at the major HSP1 TSS show signs of stuttering with non-templated adenosine at the start of the transcripts (Supplementary Figure 8c). While such stuttering has been reported previously during initiation of transcription by phage polymerases *in vitro* (38), the addition of non-templated nucleotides in human mitochondria has not been previously described.

We did not find TSS at the HSP2 position reinforcing the absence of an HSP2 promoter as indicated by previous studies (36, 37). Instead, we found three putative novel TSS across the mitochondrial genome, two on the heavy strand (positions MT:2434 and MT:3242, human GRCh38) and one on the light strand (position MT:16029). Interestingly, the novel TSS on the light strand (Supplementary Figure 8d) is located 6bp upstream for the proline tRNA and shows similar conformation to the HSP1 with stuttering of non-templated adenosine at the start of the novel transcripts. Similar nucleotide composition starting at the TSS can be found for HSP1, LSP, and the predicted novel TSS (position MT:16029) with AAAGA as the common motif.

## Discussion

In this work, we describe a novel technology, ReCappable-seq, which identifies eukaryotic TSS of all primary transcripts genome-wide at single-nucleotide resolution. Our method relies on decapping and recapping RNA with two sequential enzymatic treatments using the enzymes yDcpS and VCE, and subsequent enrichment with streptavidin. Starting with 2-5 µg of total RNA and employing standard ligation-based Illumina library kits, without the need of custom adaptor/primers, one can obtain TSS information with very low ribosomal reads content. Compared to standard CAGE protocols (e.g.N-antiCAGE) our method requires less time, is technically less difficult to perform, employs fewer steps, does not require customized reagents and should be more accessible to researchers.

ReCappable-seq identifies TSS independently of the transcribing polymerases because it captures both capped and triphosphorylated primary transcripts. In this respect, ReCappable-seq represents a significant departure from existing technologies that either target prokaryotic TSS (39, 11) or eukaryotic Pol-II TSS (17). With the growing realization of the importance of non-coding RNA, the ability of ReCappable-seq to provide a comprehensive global landscape of TSS is an important advance.

Similarly, ReCappable-seq can be applied to complex communities composed of both prokaryotic and eukaryotic organisms with the expected outcome of detecting all TSS regardless of the organism or transcribing polymerases while simultaneously depleting mature rRNA.

In this study, we focused on the human transcriptome and demonstrated that ReCappable-seq identifies TSS from transcripts derived from Pol-I, Pol-II, Pol-III, and POLRMT RNA polymerases. ReCappable-seq is in good agreement with CAGE results and other orthogonal methods for TSS identification. We further confirm the specificity of ReCappable-seq for genuine TSS by demonstrating that the TSS obtained from highly degraded samples are comparable to TSS obtained using intact RNA. Because ReCappable-seq identifies all TSS including those from highly expressed Pol-III transcripts, the sensitivity for scarce Pol-II transcripts is lower than CAGE. To achieve equivalent sensitivity as CAGE for Pol-II TSS it would be necessary to increase the sequencing depth of ReCappable-seq. Alternatively, if the focus of the work is exclusively on Pol-II transcripts, then sensitivity can be increased by removing non-Pol-II TSS with CIP pretreatment of the RNA.

ReCappable-seq can be complemented with an unenriched control library to better discriminate between primary transcripts and 5’ ends of processed/degraded RNA leading to the identification of high confidence TSS. However, these unenriched control libraries double the number of libraries to be sequenced. With an estimated <10% of false-positive TSS positions, this library is optional but is useful for studying highly expressed genes that tend to have more spurious TSS. Using the data from the ReCappable-seq CIP library allows assignment to either Pol-II or non-Pol-II TSS. Nonetheless, a definitive assignment to the transcribing polymerase may require further evidence considering the potential presence of non-canonical caps that are resistant to yDcpS or RNA structures that are CIP-resistant.

We have shown that this grouping of TSS into two distinct classes based on the sensitivity to CIP treatment is accurate for transcripts with 7mG cap structures and 5’ triphosphate ends. Interestingly, ReCappable-seq identifies some transcripts that do not have a canonical structure at their 5’ end. We show for example, that while 7SK RNA is a known Pol-III transcript, the transcripts of 7SK genes appear CIP resistant. This disparity is due to the CIP resistant gamma-methylated triphosphate found at 5’ ends of 7SK transcripts and consequently, the 7SK TSS is classified here as Pol-II consistent. The Pol-III transcribed 7SL gene, is an interesting example where ReCappable-seq identifies two adjacent TSS. The one corresponding to the annotated gene start is classified as Pol-II and is also identified by CAGE (Figure 4b). The other one located one nucleotide upstream is classified as Pol-III. Further investigation is required to uncover the exact nature of the 7SL 5’ end. ReCappable-seq recovered TSS for the major known Pol-III transcribed genes at single-nucleotide resolution with the notable exception of the U6 snRNA covered by only a few reads. It is possible that some RNA modification and/or RNA structure may affect the ability of ReCappable-seq to capture some transcripts as may be the case for U6.

Beyond the 7SL and 7SK examples, the systematic complementation of ReCappable-seq results with orthogonal datasets such as CAGE, can be used as a discovery platform to identify other interesting non-canonical capped structures. Indeed a number of non-canonical cap structures recently reported (40, 41) are expected to have distinctive outcomes. For example, the trimethyl G cap is expected to be resistant to yDcpS (13) but captured by CAGE, leading to a discrepancy between ReCappable-seq and CAGE. Despite trimethyl G cap having been described on U1, U2, U4, and U5 snRNAs (42), ReCappable-seq identifies a clear Pol-II consistent TSS at the start of these genes suggesting that perhaps only a fraction of the 5’ end of these transcripts has been fully methylated to trimethyl G.

NAD cap is another structure that will not be decapped by yDcpS (13) but captured by CAGE because of the presence of the 2’, 3’ diol on the ribose. As an example, the HSP1 TSS in the mitochondria has a strong CAGE signal consistent with the NAD cap recently reported in human mitochondria (41) while ReCappable-seq shows a CIP sensitive signal consistent with a triphosphate end. Taken together, these results suggest a mixed population (NAD caps and triphosphates) at the 5’ end of RNA transcripts which are initiated at the HSP1 TSS position.

Finally, with the ability to identify the 5’ ends of primary transcripts initiated by all the RNA polymerases, ReCappable-seq reveals a rich landscape of novel TSS that can now be studied with this new technology.

## Materials and Methods

### RNA preparation

Human lung carcinoma A549 cells (ATCC) were cultured in F12 media supplemented with L-glutamine. Total RNA was purified from cell passage number 3 using the RNeasy RNA purification kit (Qiagen Cat. No. 75142). A260:280 ratio > 2 and RIN > 9.

Human brain total RNA was obtained from Takara (Cat No. 636530). The RNA was isolated by a modified guanidinium thiocyanate method and has RIN > 9.

### ReCappable-seq procedure

Total RNA was (optionally, see text) de-phosphorylated using the Quick CIP (NEB M0525) (3 units/ µgRNA at 0.6 units/µL) and the resulting RNA was purified with the “Clean and concentrate” kit (Zymo Research R1013) using the standard protocol.

Decapping of 5 µg total RNA was performed with 200 units of yDcpS (NEB M0463S) in 10mM Bis-Tris-HCl pH 6.5, 1mM EDTA in 50 µl total volume for 1 hr at 37 °C. The de-capped RNA was purified with the “Clean and concentrate” kit as above.

Capping with 3’ Desthiobiotin GTP (DTB-GTP) was performed in 50 µl total volume with 5 µL Vaccinia capping enzyme (NEB M02080) and 0.5 mM DTB-GTP (NEB N0761), in the absence of SAM for 40 min at 37°C. The RNA was purified with “Clean and Concentrate 5, Zymo Research R1013” kit and eluted in 40 µl.

The RNA was then fragmented in 0.25X T4 Polynucleotide kinase (PNK) buffer (NEB M0201) (1 µl) at 94 °C for 2.5 minutes and immediately put on ice. The solution was further supplemented with 3 µl PNK buffer to 1X, and 1 µl of PNK (NEB M0201, 10,000 units/ml) and incubated at 37 °C for 20 minutes. The RNA was purified by AMPure beads in 50% ethanol (1 volume RNA, 2 volumes beads, 3 volumes ethanol). The material was eluted in water and a portion (20-25%) was kept as an unenriched control sample. The remaining RNA was mixed with an equal volume of hydrophilic streptavidin magnetic beads (NEB S1421) which had been pre-washed two times and resuspended in 2M NaCl, 10mM Tris-HCl pH 7.5, 1mM EDTA (SA Binding buffer). The suspension was incubated by rotation at room temperature for 1 hr. The unbound material was removed by magnetic separation and discarded. The beads containing the bound RNA were washed three times by resuspension in 300 µl of SA Binding buffer and three times in 0.25M NaCl, 10mM Tris-HCl pH 7.5, 1mM EDTA (SA Wash buffer). The bound RNA was eluted by incubation of the beads in 0.5M NaCl, 10mM Tris-HCl pH 7.5, 1mM EDTA containing 1mM D-biotin (SA elution buffer) by incubating at 37 °C for 30 minutes with occasional resuspension. The eluted material was purified with AMPure beads as above by thoroughly washing the beads and sides of the tube by washing four times with 80% Ethanol to eliminate any traces of biotin before proceeding to a second round of streptavidin binding as above.

The eluted material from the second streptavidin purification was purified with AMPure beads as above and the RNA eluted in 32 µl water. This material, in parallel with the unenriched control sample from above, were treated with 4µl RppH (NEB M0356) in 1X Thermopol buffer (NEB B90004) in a total volume of 40 µl at 37 °C for 1 hr. Afterwards RppH was deactivated by adding 0.5 µl 0.5M EDTA and incubating for 3 min at 94 °C. These samples were purified with AMPure beads as above and the RNA eluted in 15 µl of water.

This material was used for generating sequencing libraries using the Small RNA library prep Set for Illumina (Multiplex compatible) kit (NEB E7330). Typically, 11-13 cycles of PCR were used for the unenriched control RNA and 14-16 cycles for the streptavidin-enriched fraction. Indexed libraries were sequenced using Illumina Nextseq 500 (75 bp paired-end sequencing).

### RT-qPCR procedure

15 µg of total RNA from A549 cells was used for the control RT-qPCR experiments after addition of spike-in controls: ERCC mix 1 (0.1 µL per 5 µg RNA) and an in vitro transcription-generated transcript of FLuc luciferase (5 pg per 5 µg RNA). The mixture was divided into three aliquots which were subjected to the Recappable-seq procedure as described above with only one streptavidin enrichment step. One aliquot was first dephosphorylated (CIP sample), a second aliquot was not (No CIP sample) and the third was processed without the decapping step (No DcPs sample). After streptavidin enrichment all three samples were handled identically. RNA eluted from the first round of streptavidin enrichment and the corresponding unenriched control was purified with AMPure beads, and a portion corresponding to 1 µg input RNA was converted to cDNA using the LunaScript RT SuperMix cDNA synthesis kit (NEB E3010). 1% of the cDNA was used for each qPCR reaction performed with the Luna Universal qPCR Mastermix (NEB M3003) and the PCR primers listed (Supplementary Table1). The percent recovery for each target was calculated from the Ct values of enriched compared to unenriched control which was set to 100%.

### RNA-seq

The RNA-seq libraries were prepared as follows: the rRNA was depleted from 0.5 µg of A549 total RNA using the NEBNext rRNA Depletion Kit (NEB E6310). The resulting rRNA-depleted RNA was used for library construction with the NEB ultra II directional RNA-seq kit (NEB E7760). The library was sequenced on illumina Miseq (75 bp paired-end sequencing). The corresponding sequencing results are referred to as A549 RNA-seq and A549 rRNA-del RNA-seq.

### CAGE

Two replicate samples of total RNA (10 µg per sample) were sent to DNAFORM (Yokohama, Kanagawa, Japan) who performed library preparation and sequencing using the nAnT-iCAGE protocol as described in (4). First strand cDNAs were transcribed to the 5’ end of capped RNAs, attached to CAGE ‘bar code’ tags. The indexed libraries were sequenced using illumina Hiseq 2500 (50 bp single-end sequencing).

### 5’ RACE for RNA Pol-III transcripts

We determined the nucleotide sequence of transcript 5’ ends obtained from the amplified RACE products for RPPH1, RMRP and 7SL1 (Supplementary Figure 6). Total RNA from A549 cells (3 µg in 30 µl) was treated with RppH (NEB M0356) at 30 °C for 1 hr and purified by spin column. 0.5 µg of the treated RNA was used for ligation to an RNA oligonucleotide with T4 RNA ligase 1 (NEB M0204) for 1 hr at 25 °C then converted to first strand cDNA with random priming and the ProtoScript II reverse transcriptase mix (NEB E6560). Amplification reactions from this cDNA were performed with the 5’ PCR primer and a reverse primer corresponding to the target sequence using LongAmp Taq polymerase (NEB M0287). The PCR products were sequenced and the junction between the ligated oligo revealed the start sites.

Sequences of the RNA ligation oligonucleotide the 5’ forward and the reverse primers used are shown in the Supplementary Table1.

### Bioinformatics analyses

#### Datasets used for Transcription Factor (TF) binding or histone mark analysis in Figure 3 and Supplementary Figure 5

We used A549 RNA polymerase II ChIP-seq datasets (GSE33213) for polymerase binding analysis. Since A549 RNA polymerase III ChIP-seq data were not available, we used HeLa Pol-III ChIP-seq datasets (GSE20309) (14).

For TF and histone ChIP-seq, we downloaded the call sets from the ENCODE portal (34)(https://www.encodeproject.org/) with the following identifiers: ENCFF103COS ENCFF897HCS ENCFF519XGR ENCFF290AKW ENCFF331NAZ ENCFF447XHR ENCFF603JLN ENCFF538HPA ENCFF473YHH ENCFF000MYX ENCFF387NIL ENCFF721WCM ENCFF359GMW ENCFF623DEN ENCFF931GUX ENCFF000NBM ENCFF897YZE ENCFF102ZYB ENCFF648PQD ENCFF747CVL ENCFF801BLX ENCFF640ILD ENCFF322NBG ENCFF599JTK ENCFF897SFR ENCFF950NVT ENCFF172GSY ENCFF658EZL ENCFF000NDT ENCFF908FKC ENCFF137VHW ENCFF668LIG ENCFF870WJP ENCFF000NEY ENCFF454PBC ENCFF565TWZ ENCFF776IJI ENCFF785XZN ENCFF374NIY ENCFF585RGO ENCFF796BZA ENCFF512ZYH ENCFF997KRD ENCFF967MRB ENCFF829OSK ENCFF063VTN ENCFF000NIP ENCFF967EEA ENCFF539BSO ENCFF604QPX.

Bigwig files are used as it is. Bam files were compared to the appropriate control bam files using bamCompare from deeptools (version 3.3.0) (34, 43) with the following parameters --binSize 5 and --outFileFormat bigwig. Fastq files were first mapped to GRCh38 using bwa mem (44)(version 0.7.12 with default parameters) and the resulting bam files were processed as described above.

#### K562 CAGE Dataset

We used the K562 CAGE dataset generated by (17) for rRNA analysis in Supplementary Figure 2 (GSM2772305, GSM2772306 and GSM2772307, three replicates).

#### Genome and Annotations

Human genome version GRCh38/hg38 was used for all the analyses.

The comprehensive gene annotation on the primary assembly GRCh38.p5 (GENCODE v24) was downloaded from gencode: ftp://ftp.ebi.ac.uk/pub/databases/gencode/Gencode_human/release_24/gencode.v24.primary_assembly.annotation.gtf.gz

To further classify gene types, we added predicted tRNA genes derived from GENCODE v24 tRNAscan into the comprehensive gene annotation for TSS assignment in Figure 4a and Supplementary Figure 3.

The UCSC annotation used for precision and sensitivity assessment was created using the UCSC table browser and the following options: assembly: Dec2013 GRCh38/hg38; group: Genes and Gene Predictions; track: All GENCODE v24.

#### Read processing and alignment

Since CAGE reads were 50 nt, we shortened the A549 ReCappable-seq and CIP ReCappable-seq reads to retain the first 50 nt from the 5’ end. Then we processed the ReCappable-seq reads, and the paired-end RNA-seq reads by trimming the illumina adapter using Trimgalore (version 0.4.4 with parameters --length 25 --stringency 3) (https://github.com/FelixKrueger/TrimGalore). Next we aligned the trimmed reads to the human genome (GRCh38, GENCODE v24) using STAR (45) version 2.5.2 following the ENCODE data processing standard with the following parameters: --outMultimapperOrder Random, --outFilterMultimapNmax 20. For ReCappable-seq reads single end alignment was used, while for RNA-seq reads paired-end alignment was used. Therefore, local alignment with soft clipping at the 5’end was applied. Only the primary alignments, which were selected using samtools (version 1.7 with parameter -F 256) (46), were used for the TSS analysis and RNA-seq analysis.

#### rRNA analysis

To calculate the percentage of reads from processed rRNA genes, we mapped the ReCappable-seq and CAGE reads from each replicate to a human rRNA reference set (containing rRNA genes 18S, 28S, 5.8S, 12S, 16S) using bwa mem (with default parameters) and counted using samtools. We did not include the 5S rRNA genes because these genes are transcribed by Pol-III polymerase and therefore 5S is triphosphorylated.

% rRNA = (number of reads mapped to rRNA genes / number of reads mapped to human genome) x 100

#### TSS identification and analysis pipeline

The number of 5’ end tags for each position on the genome was calculated using a custom script CountTssGTF.py (https://github.com/elitaone/cappable-seq) and normalized as Tags Per Million primary mappable reads (TPM). Here, 5’ end tag refers to the 5’ most nucleotide of a read mapping to the reference genome and it corresponds to the TSS position.

To assess reproducibility, we compared the TPM for each position between the two technical replicates starting from the same RNA starting material (Supplementary Figure 1). Since the two replicates were highly correlated, in order to get more information and to compare between methods for all the ReCappable-seq and CAGE treatments, we merged the aligned reads from two replicates and randomly sampled 63 million of primary mappable reads for TSS analysis. In this study, TSS is defined as a position with TPM >= 1 (Supplementary Data 1).

To identify high confidence TSS positions (Figure 2), we compared the TPM of each TSS position obtained from the ReCappable-seq dataset with the corresponding TPM obtained from the unenriched control dataset. Positions with an enrichment Ratio >= 1 were considered as high confidence TSS (Ratio = TPM of ReCappable-seq divided by TPM of unenriched control).

#### Assignment of TSS to Pol-II or non-Pol-II

The high confidence TSS, were further classified by comparing the TPM of each TSS obtained from the untreated ReCappable-seq dataset with the corresponding TPM obtained from the CIP treated ReCappable-seq dataset. Based on the TPM Ratio (TPM untreated/TPM CIP-treated), we classified the TSS into Pol-II (TPM Ratio < 4) and non-Pol-II (TPM Ratio >= 4) (Figure 2b).

#### Assignment of TSS to annotated genes in Figure 4a and Supplementary Figure 3

To compare the defined TSS from both ReCappable-seq and CAGE to GENCODE annotated genes including tRNA prediction, for each TSS we identified the closest exon using bedtools (47) closest with the following options: -t first -D a -iu -s. We assigned a TSS to the corresponding gene if it is within 200 bp upstream or overlapping with the exon. The number of assigned unique genes or protein coding genes was counted. Non-Pol-II TSS, not associated with GENCODE annotated genes or predicted tRNAs, were clustered into individual loci where the TSS were located within 20 bp of each other.

#### Transcript body Coverage in Supplementary Figure 2b

We used a custom script (https://github.com/elitaone/transcript_body_coverage) to analyze the coverage of reads (RNA-seq) or 5’ end tags (ReCappable-seq) across the transcript body. Briefly, we calculated the number of reads or tags covering each position of transcripts longer than 300 bp. For RNA-seq we discarded low expression transcripts (FPKM < 10). For coverage analysis of ReCappable-seq and the corresponding unenriched control, we discarded transcripts whose overlapping 5’end tags contained less than n reads, where n = number of mappable reads divided by 1 million.

#### Mitochondrial TSS in Supplementary Figure 8

For TSS identification on mitochondria, we used a stringent standard to filter out the non-unique TSS positions that are from reads also mapping to nuclear chromosomal sites. More specifically, for each TSS position, we calculated the Ratio of the number of tags with MAPQ > 39 for bowtie2 and MAPQ > 3 for STAR, which indicates unique mapping respectively, to the number of total tags. Only the positions with Ratio >= 0.8 were used as TSS positions plotted on the mitochondrial genome in Supplementary Figure 8a.

#### Precision and Sensitivity assessment in Figure 3a and 3b

The assessment of precision and sensitivity was performed as described in (17) with minor modifications as shown below.

##### Reference used for precision and accuracy assessment

We mapped our high confidence A549 ReCappable-seq Pol-II TSS (Figure 2 Quadrant IV) against UCSC annotation, DNase-seq data and CAGE positions to assess precision. Only the first exon of transcripts included in the UCSC annotation were used for this comparison. The A549 DNase-seq datasets were obtained from ENCODE (bednarrow peak files: ENCFF135JRM, ENCFF698UAH, ENCFF079DJV and ENCFF045PYX, four replicates). The bednarrow peak files from all the replicates were merged for comparison with the ReCappable-seq Pol-II TSS.The A549 RNA-seq data and A549 CAGE data were generated in this study.

##### Calculation of Precision and Sensitivity

True positives (TP) were defined as the high confidence ReCappable-seq Pol-II TSS overlapping each reference including UCSC TSS annotation, CAGE positions and DNase-seq peaks as mentioned above. While False Positives (FP) were those not overlapping each reference. False negatives (FN) were defined as the UCSC TSS annotation that (1) were within any protein coding transcripts with FPKM >= cutoff as shown in Sensitivity Plot x-axis (Figure 3b) (The FPKM was quantified using RSeQC (48) based on the A549 RNA-seq data). (2) overlapped at least one A549 DNase-seq peak; (3) did not overlap any defined high confidence ReCappable-seq Pol-II TSS.

We determined the overlap using bedtools intersect using the high confidence ReCappable-seq Pol-II TSS as -a and each reference as -b with the addition of the following parameters: (1) -s option for UCSC TSS annotation comparison. (2) -s option for comparison with CAGE positions within a window of +/-1 nt.

We calculated Precision and Sensitivity as shown below:

Precision (Figure 3a) = TP / (TP + FP) *100

Sensitivity (Figure 3b) = TP / (TP + FN)

## Supplementary Data

Supplementary Data 1: List of TSS found in the human genome (GRCh38). Columns correspond to Chr (chromosome), TSS (position of the TSS, 1-coordinates), strand (+ or -), TPM (TSS tags per million), Ratio Control (TPM ReCappable-seq divided by TPM unenriched control), Ratio CIP (TPM ReCappable-seq divided by TPM CIP), Ratio CAGE ((TPM ReCappable-seq divided by TPM CAGE), feature (quadrant I, II, III or IV according to Figure 2b), source (annotation source, either GENCODE or tRNA prediction), geneType (annotation type), gene Name (name of the associated gene). N.A. corresponds to TSS of unknown origin.

## Data availability

Sequencing data and processed results were deposited and available on GEO (GSE132660). (https://www.ncbi.nlm.nih.gov/geo/query/acc.cgi?acc=GSE132660).

High confidence TSS defined by ReCappable-seq have been uploaded as a custom track in the UCSC genome browser: http://genome.ucsc.edu/s/rezo/ReCappable-seq. High confidence TSS defined by ReCappable-seq; bedGraph format; negative data values represent CIP sensitive TSS (Quadrant II), positive data values represent CIP resistant TSS (Quadrant IV, Figure 2).

## Acknowledgments

We would like to thank the NEB sequencing core facility for NGS sequencing and New England Biolabs for supporting this research.

## Notes

http://genome.ucsc.edu/s/rezo/ReCappable-seq

### Competing Interest Statement

All authors are employees of New England Biolabs Inc. a manufacturer of restriction enzymes and molecular reagents.

### Summary of Updates

Various changes in the manuscript / supplementary text

