## Supplementary Figures and Text for "ReCappable Seq: Comprehensive Determination of Transcription Start Sites derived from all RNA polymerases"

Supplementary Information

### Supplementary Figures

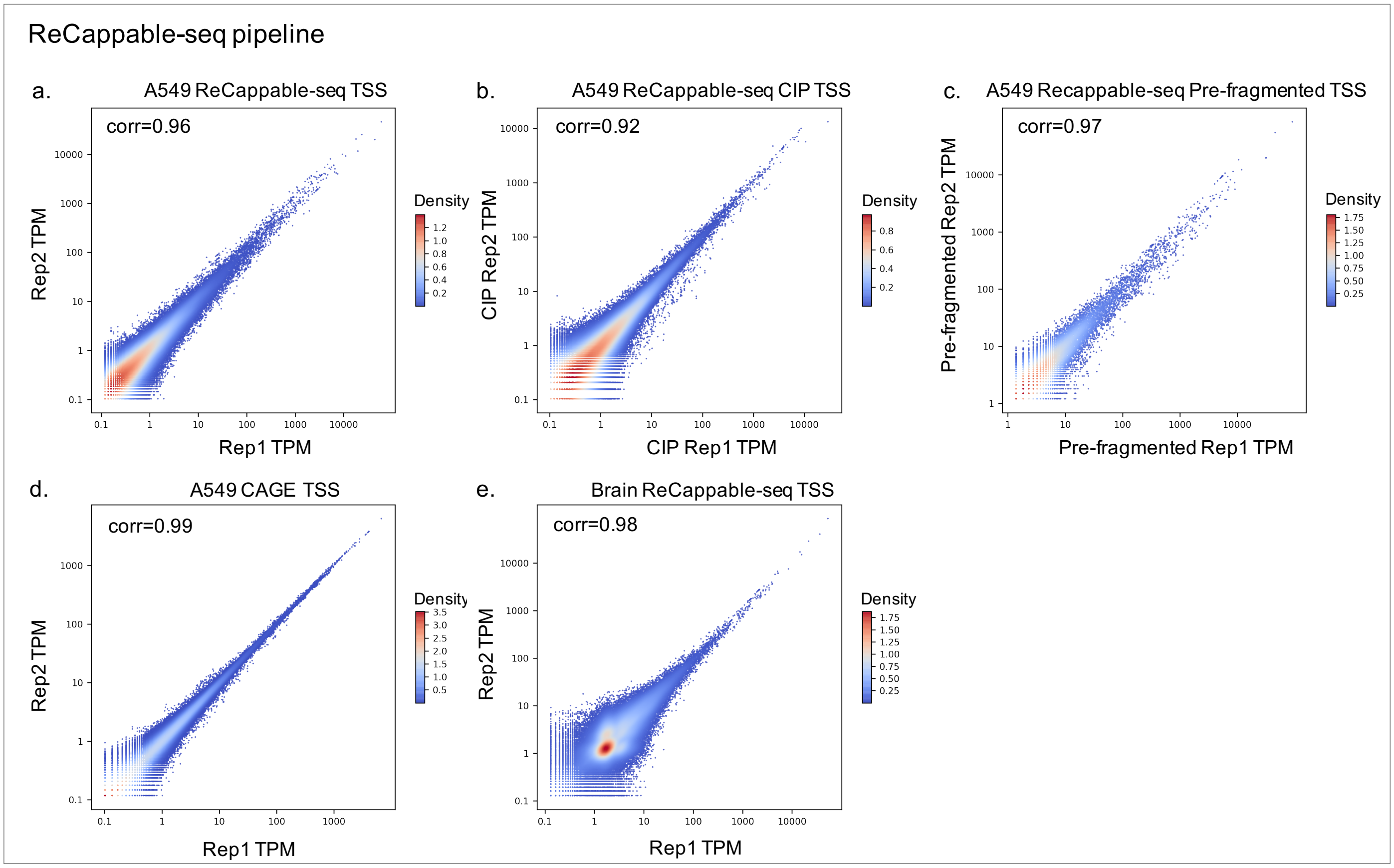


Supplementary Figure 1: Correlation between technical replicates for ReCappable-seq and CAGE experiments.

The number of reads per TSS position were normalized to the total number of mappable reads (TPM) for both technical replicates. Positions with TPM >= 1 were considered. Replicate analysis for ReCappable-seq performed on A549 intact RNA (a), CIP + ReCappable-seq performed on A549 intact RNA (b), ReCappable-seq performed on pre-fragmented A549 RNA (c), CAGE experiments on A549 intact RNA (d) and ReCappable-seq performed on brain total RNA (e).

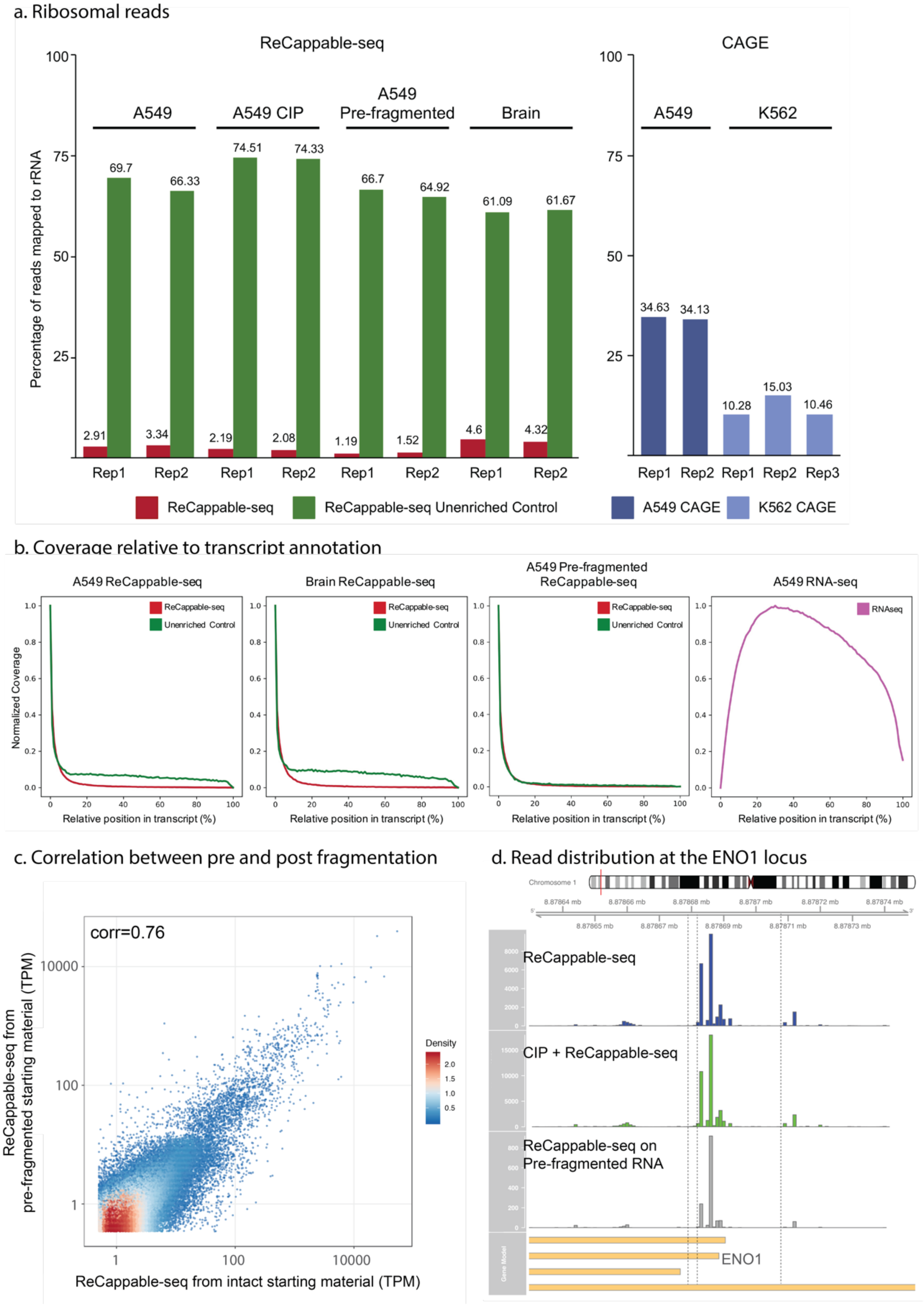


Supplementary Figure 2:

a. Percentage of mapped reads mapping to ribosomal loci in the ReCappable-seq and the unenriched control datasets. Unenriched control datasets (green) denote ReCappable-seq datasets for which the streptavidin step has been omitted. b. Read profiles relative to position in the annotated gene bodies (transcript body coverage) for the A549 ReCappable-seq, ReCappable-seq performed on pre-fragmented A549 RNA, the brain ReCappable-seq and A549 RNA-seq. For the

ReCappable-seq libraries (red) the unenriched control (minus streptavidin) libraries are also shown (green). c. Correlation (corr = 0.76, P-value < 2.2e-16) between TPM of ReCappable-seq derived from intact starting material (x axis) and pre-fragmented starting material (y axis). d. Mapping profiles at single nucleotide resolution of ReCappable-seq (top panel), CIP + ReCappable-seq (middle panel) and ReCappable-seq performed on pre-fragmented RNA (bottom panel) upstream of the ENO1 (Enolase 1) gene. Note that the reads and the gene are located in the reverse strand.

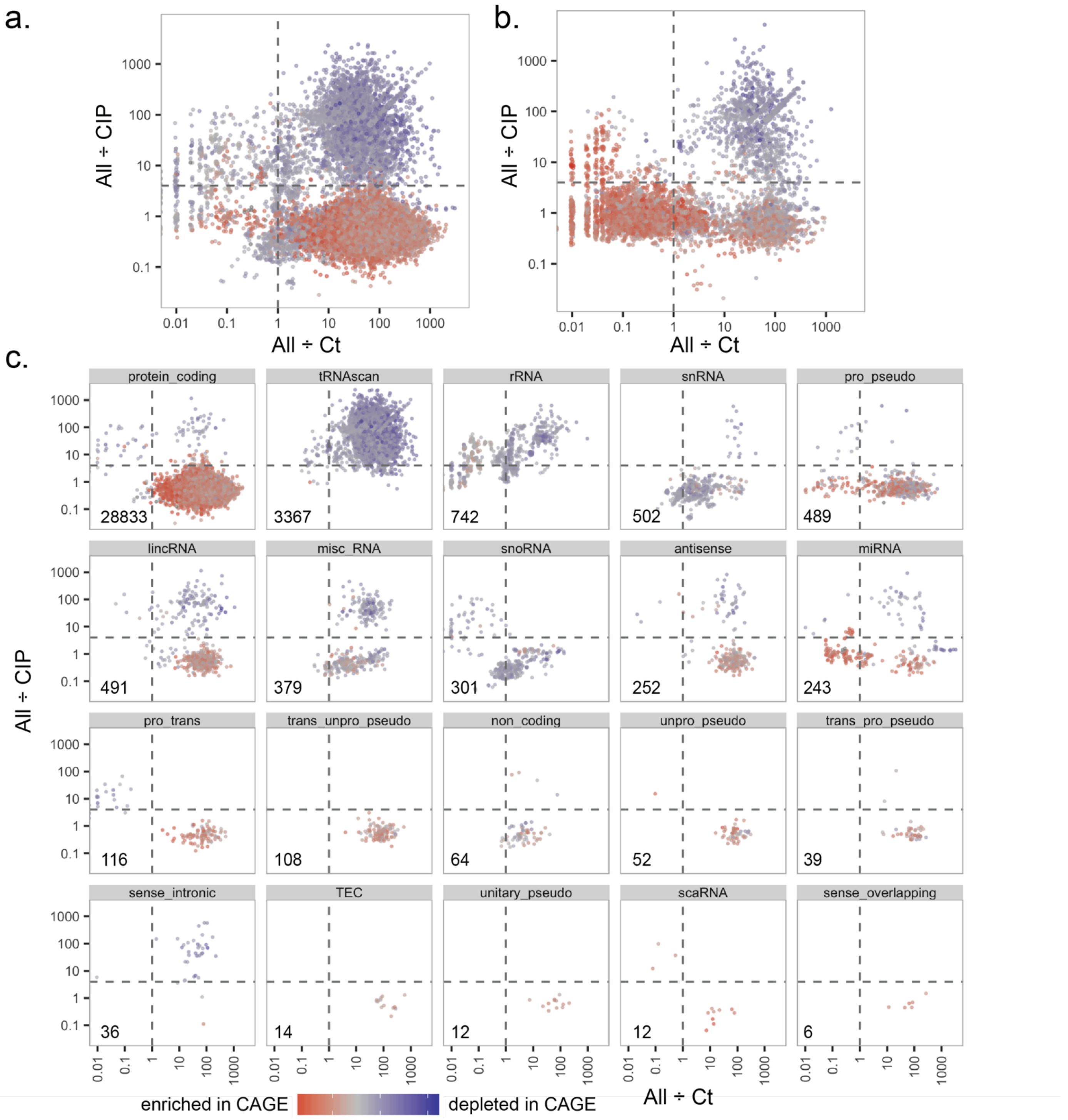


Supplementary Figure 3:

Distribution of candidate TSS derived from ReCappable-seq based on the ratio between TPM-ALL and TPM-Ct (x axis) and the ratio between TPM-ALL and TPM-CIP (y axis) plotted as in Figure 2. Color of the candidate TSS dots represents the ratio between TPM-ALL and TPM-CAGE. a. Subset of the candidate TSS (36,095) assigned to genes in GENCODE annotation including tRNA prediction. b. Subset of the candidate TSS (6,893) not assigned to any gene. c. All the candidate TSS positions shown in (a) assigned to the indicated gene types (GENCODE and tRNA prediction). The number on the left-bottom corner of each box represents the number of candidate TSS (positions shown in all four quadrants) assigned to the corresponding gene type.

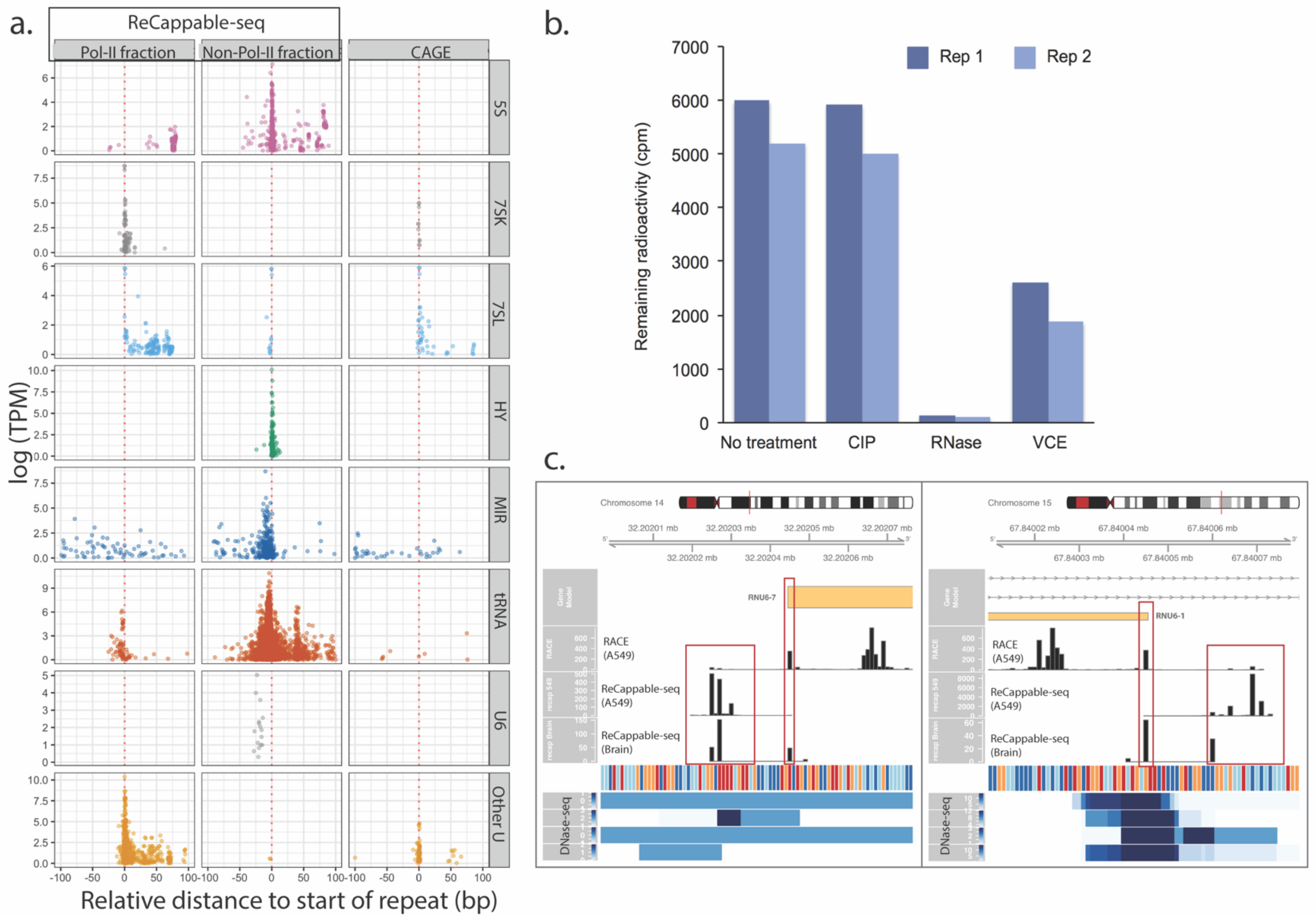


Supplementary Figure 4:

a. Position of the Pol-II consistent TSS (left panels), non-Pol-II consistent TSS (middle panels) and CAGE TSS (right panel) relative to start sites for all 5S, 7SK, 7SL, HY, MIR, tRNA, U6 and other snoRNA genes in A549. Annotation of repeats are derived from UCSC repeat masker. b. Remaining acid insoluble radioactivity (cpm, y-axis) after TCA precipitation of a 300 nucleotide RNA made by *in vitro* transcription which had been labeled with a tritiated methyl group on its 5' gamma phosphate (using MePCE) and incubated with either no addition (values=5992, 5206 cpm) CIP (5916, 5003 cpm), VCE (2619, 1874 cpm) or RNASE I (139, 97 cpm) (Materials and Methods). c. RACE and ReCappable-seq TSS results obtained for two U6 loci. The four bottom tracks correspond to read density from ENCODE DNase-seq performed on A549 cells (ENCFF473YHH, ENCFF809KIH, ENCFF821UUL, ENCFF961WXW). The mapping was done using STAR with parameters --outFilterMultimapNmax 500 --winAnchorMultimapNmax 500, which allow for multiple mapping sites.

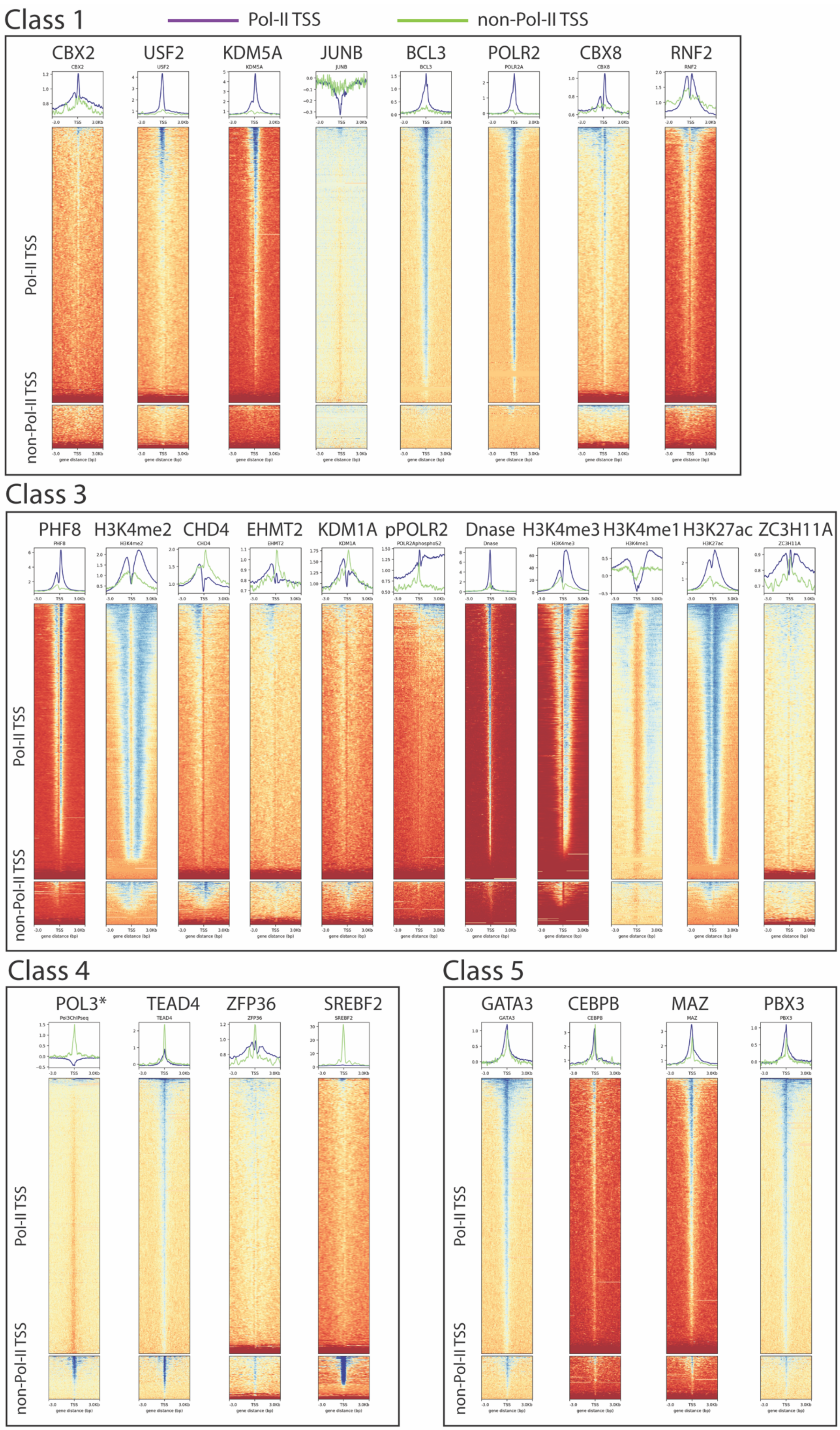

Class 1
Pol-II TSS
non-Pol-II TSS
CBX2
USF2
KDM5A
JUNB
BCL3
POLR2
CBX8
RNF2
Class 3
PHF8
H3K4me2
CHD4
EHMT2
KDM1A
pPOLR2
Dnase
H3K4me3
H3K4me1
H3K27ac
ZC3H11A
Class 4
POL3*
TEAD4
ZFP36
SREBF2
Class 5
GATA3
CEBPB
MAZ
PBX3

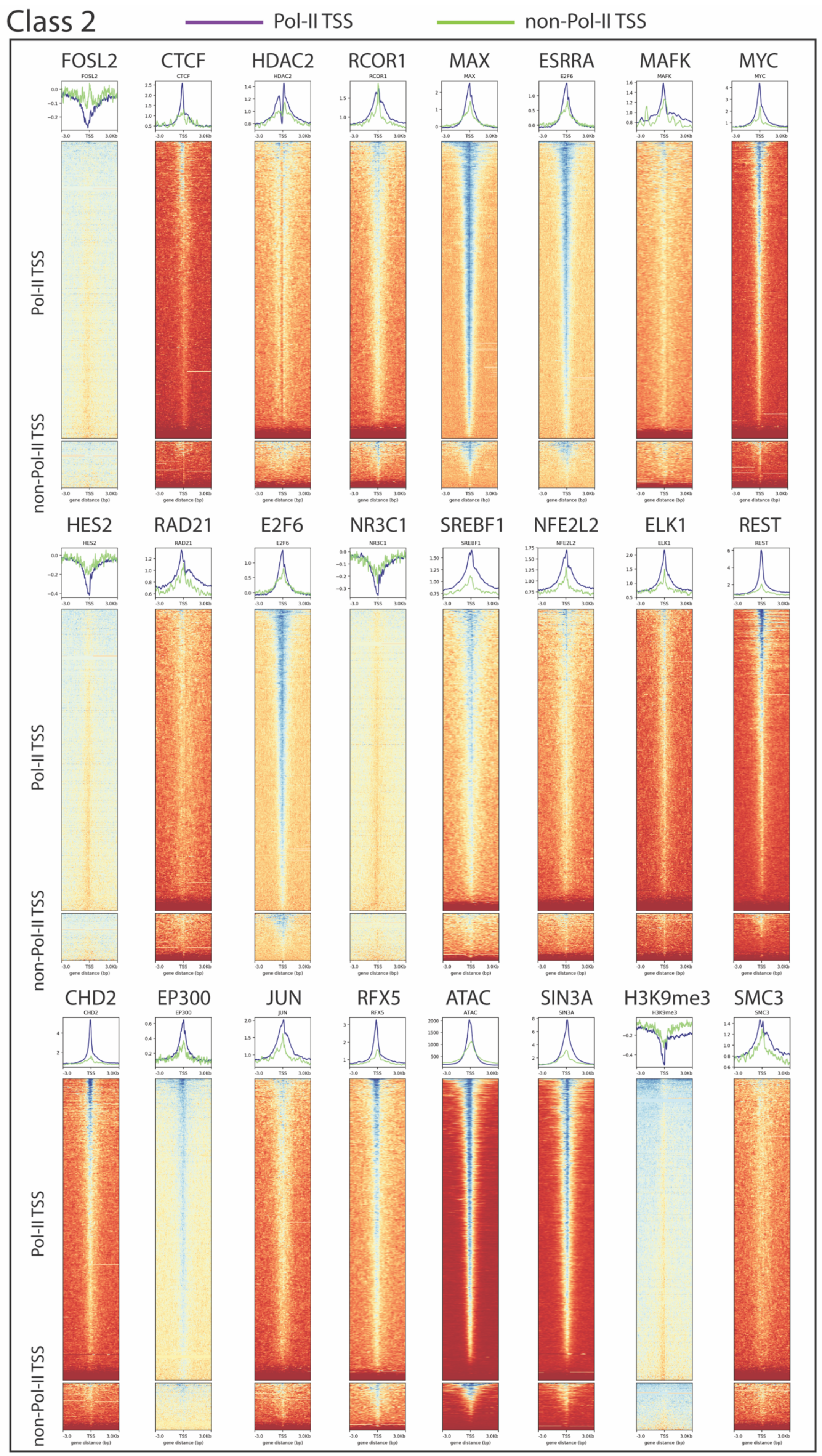

Class 2
Pol-II TSS
non-Pol-II TSS
FOSL2
CTCF
HDAC2
RCOR1
MAX
ESRRA
MAFK
MYC
HES2
RAD21
E2F6
NR3C1
SREBF1
NFE2L2
ELK1
REST
CHD2
EP300
JUN
RFX5
ATAC
SIN3A
H3K9me3
SMC3
Pol-II TSS
non-Pol-II TSS

Supplementary Figure 5:

ENCODE ChIP-seq and chromatin profiles plotted in a 6 kb window centered at the Pol-II (purple) and non-Pol-II (green) TSS derived from ReCappable-seq. Epigenetic marks were derived from ENCODE data for A549 cells except for the Pol-III binding sites (POL3*) that were obtained from Hela cells [(1)]. Profiles of epigenetic marks and DNA binding proteins were classified into 5 classes according to their distinctive profile on Pol-II-consistent TSS compared to non-Pol-II-consistent TSS. Class 1 corresponds to marks that have a distinct profile on Pol-II consistent TSS but show no distinct profile on non-Pol-II consistent TSS. Class2 corresponds to marks that have the same profile on Pol-II and non-Pol-II consistent TSS but with a weaker intensity for non-Pol-II consistent TSS. Class 3 corresponds to marks that have a distinct profile on Pol-II compared to non-Pol-II consistent TSS. Most of the histone marks fall into this category. Class 4 corresponds to marks that have a strong distinct profile on non-Pol-II consistent TSS but show no or weaker or “opposite” profile on Pol-II consistent TSS. Finally, class 5 corresponds to marks that have the same distinct profile on Pol-II and non-Pol-II consistent TSS.

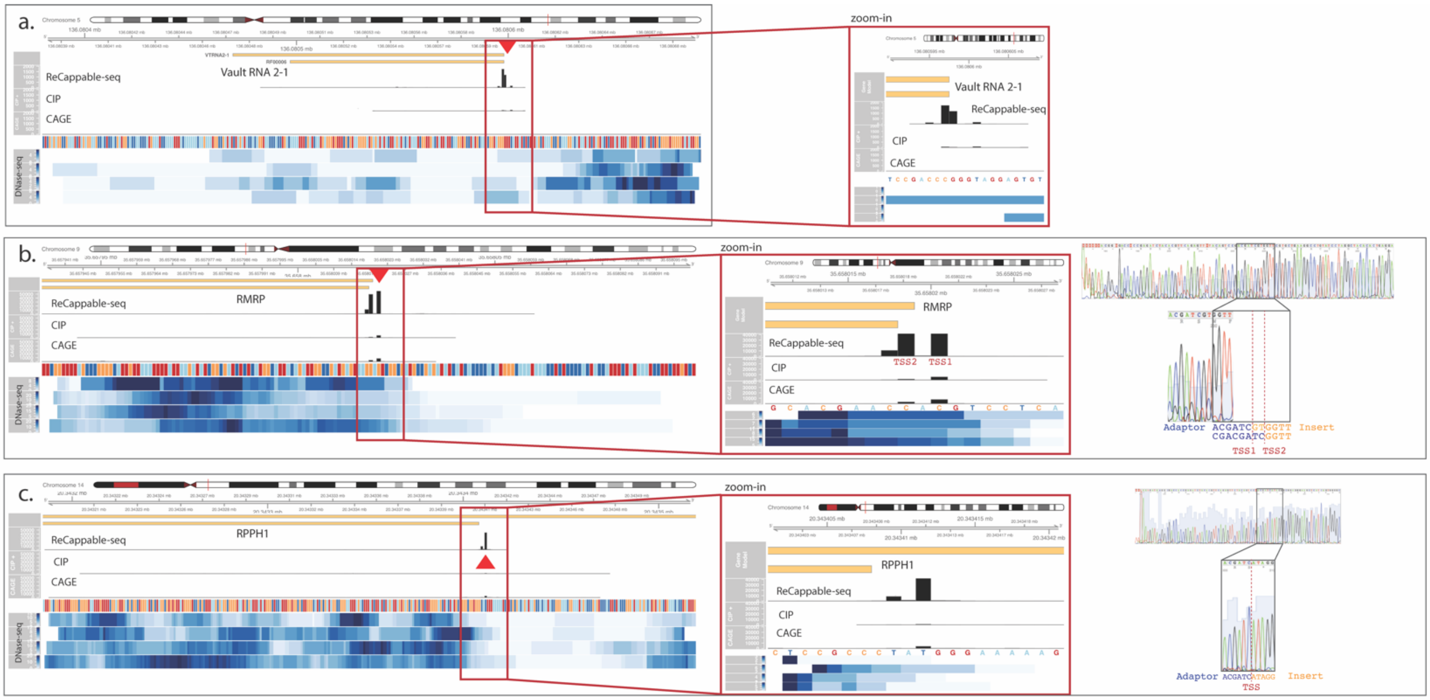


Supplementary Figure 6:

Genomic regions of Vault 2.1 gene (a), RMRP gene (b), and RPPH1 gene (c). Red arrowheads denote the positions with the highest density of reads starting at these locations. RACE experiments (Sanger traces of RT-PCR products, right panels) were performed for the RMRP and the RPPH1 transcripts revealing one transcript start for RPPH1 and two transcript starts for RMRP.

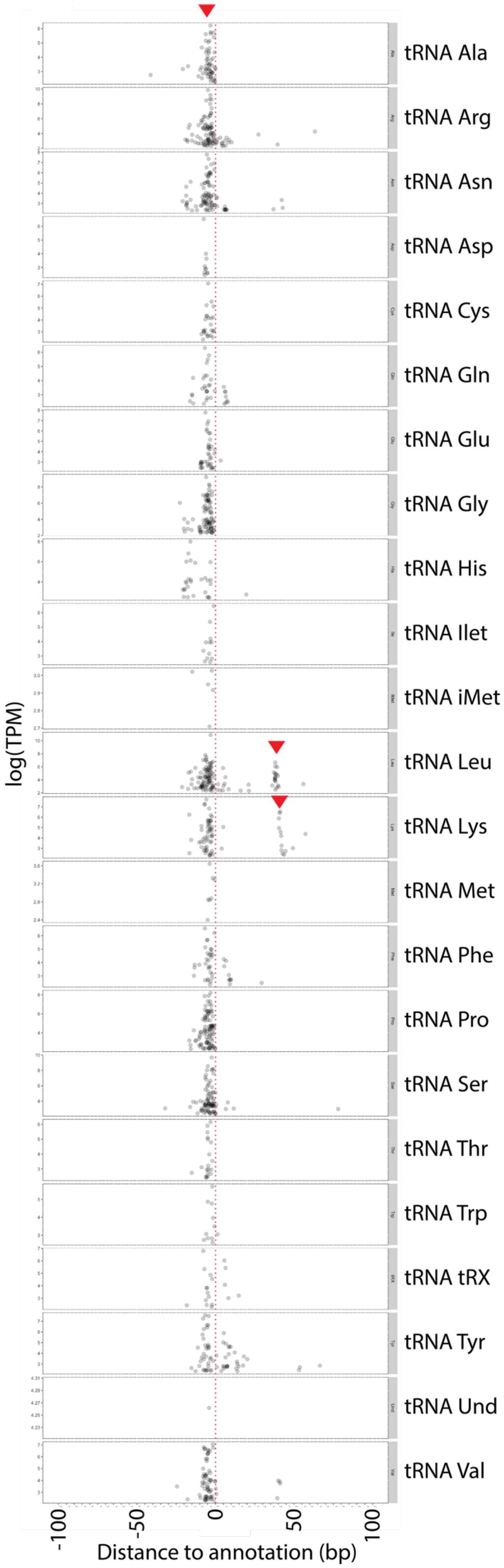
tRNA Ala
tRNA Arg
tRNA Asn
tRNA Asp
tRNA Cys
tRNA Gln
tRNA Glu
tRNA Gly
tRNA His
tRNA Ilet
tRNA iMet
tRNA Leu
tRNA Lys
tRNA Met
tRNA Phe
tRNA Pro
tRNA Ser
tRNA Thr
tRNA Trp
tRNA tRX
tRNA Tyr
tRNA Und
tRNA Val
log(TPM)
-100
-50
0
50
100
Distance to annotation (bp)

Supplementary Figure 7: Profiles of ReCappable-seq TSS around tRNA.

Non-Pol-II TSS distribution around mature tRNA annotation where 0 bp is the 5' nucleotide of the mature RNA. tRNAs are classified according to the type of tRNA. Red arrowheads denote unexpected TSS clusters. Y-axis represents the strength of the TSS (in log10(TPM)).

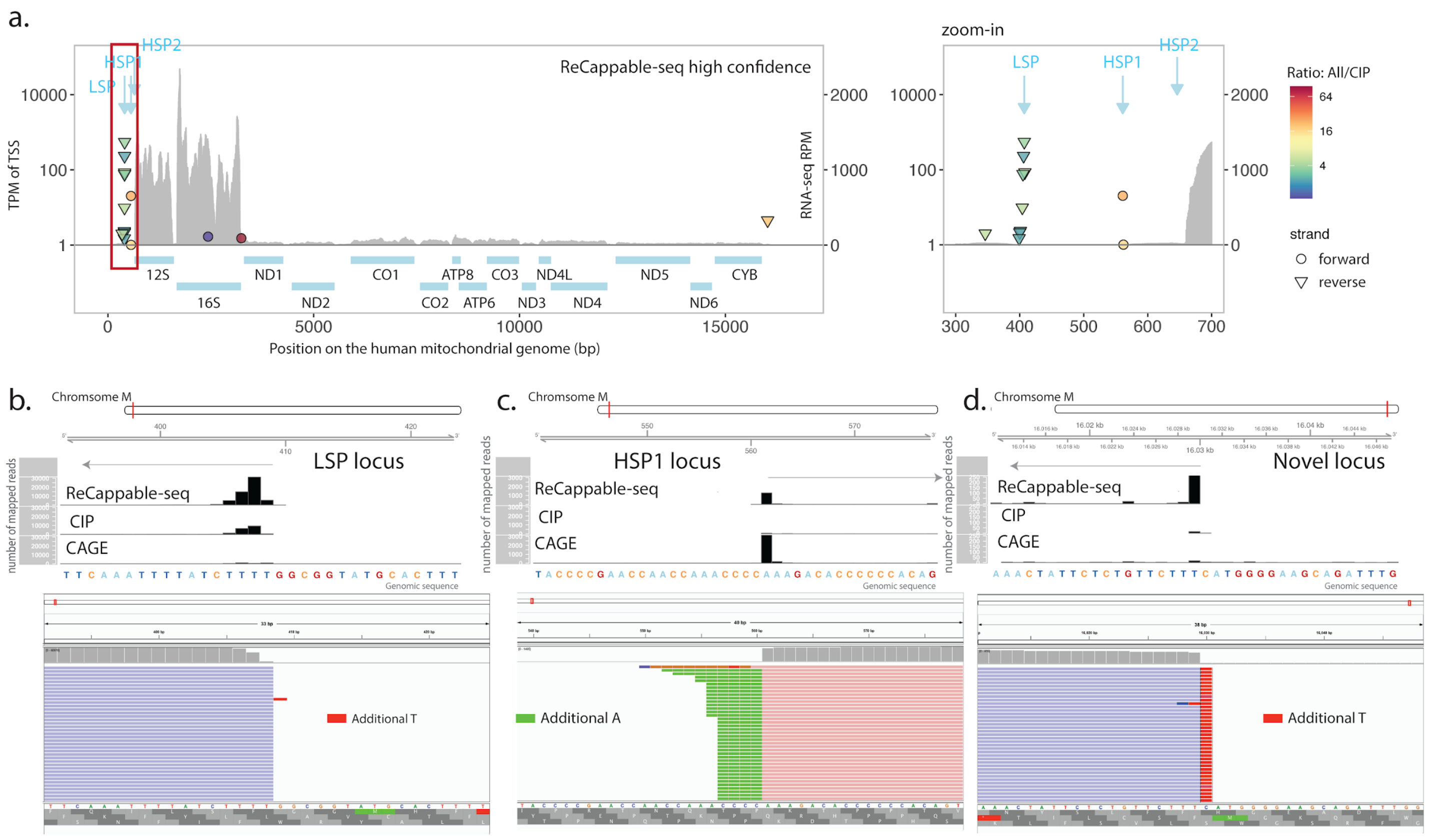


Supplementary Figure 8: Identification of mitochondrial TSS based on STAR mapping.

a. Positions of TSS detected on the mitochondrial genome in the forward (circles) and reverse (triangles) orientation. Color denotes the ratio between ALL and CIP (as shown in Figure 2). Positions of the known TSS are indicated with blue arrows. Gene expression measured by RNA-seq (grey) is calculated in RPM (Reads per million of mappable reads). Only the uniquely mapped TSS positions are shown (See Materials and Methods). Red rectangle marks the zoomed area shown on the right. b. LSP (reverse strand) locus, c. HSP1 (forward strand) locus, d. Locus containing a putative novel light strand TSS. Upper panels show the density of reads starting at various genomic locations in the mitochondrial genome for ReCappable-seq (top), CIP (middle) and CAGE (bottom). The lower panel displays an IGV rendering of a subset of the mapped reads in the forward (pink) and reverse (blue) orientation. Soft-clipped sections of the reads are colored according to the nucleotide type: A (green), T (red), G (orange) and C (dark blue).

### Supplementary Text 1: Comparison with publicly available ChIP-seq data

We classified ChIP-seq data into 5 classes according to their profiles at Pol-II and non-Po-lII TSS : [1] distinctive profile observed only at Pol-II TSS, [2] same distinctive profile but weaker signal at non-Pol-II TSS, [3] distinctive but different profiles at Pol-II and non-Pol-II TSS, [4] Distinctive profiles at non-Pol-II TSS and no (or weaker or opposite) profiles at Pol-II TSS and [5] same profile at Pol-II and non-Pol-II TSS (Supplementary Figure 5).

Most of the chromatin landscape of Pol-III-transcribed genes resembles that of Pol-II-transcribed genes in agreement with the literature [2, 3]. Nonetheless, with the ability to precisely position epigenetic marks relative to TSS, subtle differences in the profile of these marks can be highlighted. For example, H3K4me3 mark is present in both Pol-II and non-Pol-II TSS (Supplementary Figure 5). Nonetheless, the profile of this histone mark is different: while Pol-II TSS have H3K4me3 marks on both 3' and 5' of the TSS, H3K4me3 is mainly present 5' upstream of non-Pol-II TSS.

Interestingly we find examples of specific transcription factor binding profiles at non-Pol-II genes that have not been previously reported. For example, SREBP2 (coding by SREBF2 gene) is a ubiquitously expressed transcription factor known to control cholesterol homeostasis by stimulating transcription of sterol-regulated genes [4]. We found that SREBP2 is binding almost exclusively at non-Pol-II TSS (class 4, Supplementary Figure 5). This result strongly suggests a role of SREBP2 in regulating non-Pol-II genes.

### Supplementary Methods

#### Generation of gamma 3H-methyl triphosphate 300 nt long RNA transcript

A MePCE enzyme preparation was obtained as follows: the Human MePCE gene (NCBI ref #NP_062552.2 coding for a 689 amino acid protein) was expressed in *E. coli* with a His tag and purified by IMAC chromatography. 530 picomoles of a 300 nucleotide RNA *in vitro* transcript was incubated in a 200 µl reaction containing 20mM Tris pH 8.0, 0.5mM DTT, 600 picomoles of tritiated SAM, 2mM EDTA, 50mM KCl and 5% glycerol for 2.5 hours at 30 ºC with 20 µl of MePCE enzyme preparation. The RNA was isolated from the reaction by addition of 2 volumes of AMPure beads, washed 2X with 80% ethanol and eluted with 50 µl 1mM Tris pH 7.5 and 0.1mM EDTA.

#### Gamma methyl phosphate is CIP resistant

A 300 nt RNA made by *in vitro* transcription which had been labeled with a tritiated methyl group on its 5' gamma phosphate was incubated with either CIP, VCE or RNASE I, subjected to TCA precipitation and the remaining acid insoluble radioactivity was determined. Treatment with CIP does not decrease the insoluble radioactivity whereas the majority of the radioactivity is removed by the triphosphatase activity of VCE or RNASE (Supplementary Figure 4b). This demonstrates that the gamma phosphate when methylated is resistant CIP while it can be removed by VCE, suggesting that gamma methyl-triphosphate RNA should be subject to capping with VCE. 5 picomoles of gamma methyl-triphosphate RNA was treated with 1 µl of the following enzymes in 10 µl parallel reactions in the corresponding buffers for 60 minutes at 37 ºC; calf intestinal phosphatase (CIP, NEB M0525) in CutSmart buffer (NEB B7204); Vaccinia capping enzyme (VCE, NEB M02080) and no treatment sample in VCE buffer and RNASE If (NEB M0243) in NEB buffer 3 (NEB B7003). The

RNA was TCA-precipitated onto 3M Whatman paper discs and washed 4X with 5% TCA, washed 2X with ethanol, dried and tritium radioactivity counts determined by scintillation.

##### 5' RACE for U6 transcripts

Total RNA from A549 cells (2 µg) was treated with 12.5 units of RppH (NEB M0356) in 1X Thermopol buffer (NEB B9004) in a 25 µl reaction volume at 37 ºC for 1 hr, followed by addition of 0.1 µl of Proteinase K (NEB P8107) at 37 ºC for 5 minutes. The RNA was purified with the "Clean and concentrate" kit (Zymo Research R1013) using the standard protocol. The resulting RNA was ligated with the NEB 5' SR Adaptor (NEB E7328) by first dissolving the 5' SR Adaptor in 120 µl of water, adding one µl of the adaptor to 1 µg of RNA in a total volume of 13 µl and heating to 75 ºC for 5' and quick cool on ice. 13 µl of RNA/adaptor was incubated in 1X RNL1 buffer, 1 mM ATP, 15% PEG 8000, and 1.2 µl NEB T4 RNL1 ligase (M0437M) in a total reaction volume of 25 µl for 3.5 hours at room temperature. 5 µl of the ligation reaction was incubated in a 50 µl reverse transcriptase reaction with 500 units of Protoscript II (NEB M0368), 1 µM U6 primer 2R for 15 minutes at 48 ºC. The reaction was then incubated with 100 units RNase If (NEB M0243) for 10 minutes at 37 ºC, and then 20 minutes at 70 ºC. The cDNA was amplified for 30 cycles with the 5' NEB SR Primer and U6 2R primer using the LongAmp Taq polymerase (NEB M0287).

The amplified product was electrophoresed on a 1.5 % agarose gel and the region of the gel corresponding to a range of 100 to 250 nucleotides was excised from the gel and the cDNA extracted from the gel. The cDNA was amplified with SR primer and U6 3 reverse primer containing the illumina adapter sequence for 4 cycles, and then further amplified with illumina index primer. The cDNA was sequenced on an illumina Miseq (150 bp paired-end sequencing).
